# Transcriptional reprogramming and microbiome dynamics in garden pea exposed to high pH stress during vegetative stage

**DOI:** 10.1101/2024.10.05.616821

**Authors:** Asha Thapa, Md Rokibul Hasan, Ahmad H. Kabir

## Abstract

High soil pH severely impacts plant growth and productivity, yet the transcriptomic changes and microbial dynamics underlying stress adaptation in garden pea (*Pisum sativum* ssp. *hortense*) remain unclear. This study demonstrates that high soil pH leads to stunted growth, reduced biomass, impaired photosynthesis, and nutrient status in garden pea. Further, disruption in key nitrogen-fixing bacteria (*Rhizobium indicum, R. leguminosarum,* and *R. redzepovicii*), along with the downregulation of *NifA* and *NifD* genes and upregulation of *NifH* in nodules highlight the critical role of micronutrient balance in legume-microbe symbiosis and a compensatory response to maintain nitrogen status. RNA-seq analysis revealed extensive transcriptional reprogramming in roots, characterized by the upregulation of oxidative stress response genes (e.g., oxidoreductase and glutathione transferase activities, metal ion transporters) and the downregulation of genes related to ammonia-lyase activity and ion binding, reflecting broader disruptions in nutrient homeostasis. KEGG pathway analysis identified enrichment of MAPK signaling pathway, likely interacting with other pathways associated with stress tolerance, metabolic adjustment, and structural reorganization as part of adaptive responses to high pH. Root microbiome analysis showed significant enrichment of *Variovorax, Shinella,* and *Chaetomium*, suggesting host-driven recruitment under high pH stress. Stable genera such as *Pseudomonas, Novosphingobium, Mycobacterium, Herbaspirillum,* and *Paecilomyces* displayed resilience to stress conditions, potentially forming core microbiome components for adaptation to high pH. In a targeted study, inoculation of plants with an enriched microbiome, particularly *C. globosum*, under high pH conditions improved growth parameters and increased the abundance of *Stenotrophomonas* and *Pseudomonas* in the roots. It suggests that these bacterial genera may act as helper microbes to *C. globosum*, collectively promoting stress resilience in pea suffering from high pH. These findings provide a foundation for microbiome-aided breeding programs and the development of microbial consortia to enhance the adaptation of pea plants to high pH conditions.

## 1. Introduction

Soil pH is a critical factor influencing plant growth and development. High pH, or alkaline conditions, can significantly impact plant growth and micronutrient availability (Xia et al. 2024). The elevated levels of Na^+^, HCO^−3^, and CO ^−3^ in soils contribute to an increase in soil pH (Li et al. 2018). In alkaline soils, the solubility of many essential micronutrients, such as iron, manganese, zinc, and copper decreases. These micronutrients act as necessary components for various enzymes and proteins, which are pivotal for numerous physiological functions such as photosynthesis, protein synthesis, nitrogen metabolism, and respiration (Kaya et al. 2019). The amount of micronutrients, especially iron that plants can access in the soil is insufficient for their growth requirements, particularly in high pH soil conditions (Zhu et al. 2019). High pH conditions primarily manifest in plants through symptoms such as chlorosis, stunted growth, necrosis, distorted growth patterns, root dysfunction, increased susceptibility to diseases, and altered coloration, all of which are linked to impaired nutrient uptake, particularly of micronutrients like Fe, zinc, and manganese (Turner et al. 2020; Samborska-Skutnik et al. 2020). Thus, alkalinity-induced micronutrient deficiency can greatly hamper the structure and operation of photosynthetic machinery in plants (Singh et al. 2021).

Plants cope with soil alkalinity through various strategies that enhance their resilience and nutrient uptake. They often release organic acids from their roots to lower the pH around the root zone, improving the availability of essential nutrients (Barrow and Hartemink 2023; Ma et al. 2022). Physiologically, plants employ specialized ion transport mechanisms to uptake beneficial cations, and they may regulate their internal pH using proton pumps (Ku et al. 2022; Kabir et al. 2012). Morphologically, some species develop extensive root systems to access deeper soil layers and establish mycorrhizal associations, enhancing nutrient absorption (Zhang et al. 2020; Chen et al. 2016). Particularly in high pH-induced iron deficiency, dicot plants are characterized by an increase in the activity of a plasma membrane-bound reductase, which leads to higher rates of Fe(III) reduction and the subsequent breakdown of Fe(III)-chelates at the plasma membrane (Kabir et al. 2012; Marschner et al. 1986). Plant roots may release reducing and/or chelating substances (e.g., phenolics) into the rhizosphere, further mobilizing sparingly soluble micronutrients under soil alkalinity (Wu et al. 2012). Furthermore, plants utilize a variety of genes to cope with soil alkalinity, facilitating their adaptation to high pH conditions. Ion transporter genes, including H^+^-ATPases and Na^+^/H^+^ antiporters, help maintain ion homeostasis by regulating the uptake of essential cations while preventing toxic sodium accumulation (Assaha et al. 2017). Additionally, ferric reductase genes like *FRO2* are upregulated to reduce Fe(III) to Fe(II), aiding Fe uptake under high soil pH (Kabir et al. 2012).

In legume plants, the nodulation process initiates through a symbiotic interaction with nitrogen-fixing bacteria, leading to the formation of root nodules. These structures are crucial as they house the nitrogenase enzyme responsible for converting atmospheric nitrogen into a usable form for plants (Wang et al. 2018). High pH can impair the formation and function of these nodules, leading to reduced nitrogen fixation and overall plant health (Sankari et al. 2022; Oláh et al. 2001). The availability of iron plays a crucial role in the biosynthesis of several proteins essential for nodulation (Brear et al. 2013; Abadía, 2000). Nodulation bacteria, primarily rhizobia, play a crucial role in the nitrogen fixation process in legume crops (Shumilina et al. 2023; Dusha 2002). Studies have revealed that insufficient iron levels due to soil alkalinity can hinder both the growth of legume crops and their capacity for symbiotic nitrogen fixation (Slatni et al. 2011). Furthermore, high pH can lead to increased competition from symbiotic bacteria and changes in the microbial community structure in the rhizosphere (Deng et al. 2024). Although limited research has been conducted on the dynamics of bacterial communities in plants exposed to high pH, carbonate ions have been shown to significantly affect rhizosphere fungal communities (Luo et al. 2023).

Garden pea (*Pisum sativum* ssp. hortense) is an important source of micronutrients for both humans and animals (Dahl et al. 2012). However, high soil pH in peas results in significant yield loss due to impaired photosynthetic efficiency, leading to chlorosis and stunted growth (Meisrimler et al. 2016). While previous studies have explored plant responses to high pH stress (Qiao et al. 2024; Turner et al. 2020, they predominantly focus on *in vitro* or hydroponic conditions, leaving a significant gap in understanding how plants navigate this stress under more realistic and mature growth scenarios. Moreover, the impact of high pH on the transcriptional response in the roots of garden pea during vegetative growth phase remains unclear. Specially, little is known about how root-associated microbial communities respond to such conditions and how they can be leveraged to mitigate high pH stress. Therefore, this multi-omics study aims to investigate the transcriptional shift and microbial dynamics in the nodules and roots of pea plants during vegetative growth phase subjected to high pH. We also investigated whether the inoculation of an enriched microbiome, recruited by host determinants, can induce resilience to garden pea exposed to high pH in soil.

## 2. Materials and methods

### 2.1. Plant cultivation and growth conditions

Garden pea seeds (var. Sugar Snap) were surface sterilized by immersing them in a 4% sodium hypochlorite solution for 5 min, followed by three thorough rinses with sterile water. The sterilized seeds were then placed in trays and incubated in the dark at 25°C for two days to promote germination. After three days, uniformly germinated, healthy seedlings were transferred into 500 g soil pots containing an equal blend of natural field soil and commercial potting mix (1:1 ratio), for two distinct treatments: control (soil without lime, pH ∼6.5) and high pH (soil amended with 5.0 g NaHCO_3_ and 3.5 g CaCO_3_, pH ∼8.0). Throughout the cultivation period, 15 mM NaHCO_3_ was added weekly to maintain its pH to 8.0 (Kabir et al. 2012). Furthermore, the effect of different microbial consortia on the growth responses of plants was studied in different combinations of *Chaetomium globosum* (Carolina Science, North Carolina) and *Variovorax paradoxus* (NRRL: B-1908, ARS Culture Collection, USA) inoculated once in a week at a dose of 1 ×10 cfu per mL. Pea seedlings were cultivated in seven different conditions: untreated controls, high pH, high pH + *C. globosum*, high pH + *V. paradoxus*, high pH + *C. globosum* + *V. paradoxus*, *C. globosum* +, *V. paradoxus* and *C. globosum* + *V. paradoxus*. The physicochemical characteristics and nutritional status of the soil are detailed in Supplementary Table S1. The experiment followed a randomized complete block design with nine replications per treatment. Plants were maintained in a growth chamber set to a photoperiod of 10 hours of light and 14 hours of darkness (250 μmol m ² s ¹) at an approximate temperature of 25°C for five weeks. Irrigation was carried out every other day using deionized water to ensure consistent hydration.

### 2.2. Measurement of morphological features and chlorophyll fluorescence kinetics

The shoot height and the longest root of each plant were measured using a measuring tape. Harvested shoots and roots were dried in an electric oven at 70°C for 3 days, and their dry weights were recorded. Roots were gently washed to remove adhering soil particles, and nodules were carefully harvested, counted using a magnifying lens, and stored at –80°C for further analysis. Additionally, chlorophyll score and chlorophyll fluorescence kinetics (OJIP) measurements, including Fv/Fm (maximal photochemical efficiency of PSII) and Pi_ABS (photosynthetic performance index) were evaluated on the youngest fully expanded leaves. Chlorophyll content was measured with a handheld chlorophyll meter (AMTAST, United States), and fluorescence measurements were taken with a FluorPen FP 110 (Photon Systems Instruments, Czech Republic) after dark-acclimating the leaves for 1 hour prior to data collection.

### 2.3. Nutrient analysis in root and shoot

Nutrient analysis was performed on both root and young leaf samples. Roots were carefully detached, rinsed under running tap water, and immersed in a 0.1 mM CaSO_4_ solution for 10 min before being rinsed with deionized water. Young leaves were washed separately with deionized water to remove any surface impurities. The cleaned samples were then placed in envelopes and dried in an electric oven at 75°C for three days. Elemental concentrations were analyzed using inductively coupled plasma mass spectrometry (ICP-MS) at the Agricultural & Environmental Services Laboratories, University of Georgia. The concentrations were quantified by comparing ion peak intensities in the spectrum to calibration standards.

### 2.4. RNA-sequencing and bioinformatics analysis

RNA-seq analysis was performed on root samples. To ensure the removal of surface contaminants, roots were thoroughly cleaned by washing twice in sterile phosphate-buffered saline with 10 seconds of vortexing, followed by two rinses with sterile water. The cleaned roots were ground into a fine powder using a pre-cooled pestle and mortar with liquid nitrogen. Total RNA was extracted from the powdered material using the SV Total RNA Isolation System following the manufacturer’s instructions (Promega, USA). For each sample, 1 μg of RNA with an RIN value exceeding 8 was used for RNA-seq library preparation. Libraries were prepared using the KAPA HiFi HotStart Library Amplification Kit (Kapa Biosystems, USA) and sequenced on an Illumina NovaSeq 6000 platform with 0.2 Shared S4 150 bp paired-end sequencing at the RTSF Genomics Core, Michigan State University. The sequencing produced high-quality data, with 92.5% of reads passing filters and achieving a quality score of ≥ Q30.

Bioinformatics analysis of raw FastQ files was conducted using Partek Flow genomic analysis software (Partek, St. Louis, MO, USA). Adapter sequences and low-quality reads were removed using Trimmomatic (Bolger et al.2014), resulting in high-quality clean reads. The garden pea reference genome was downloaded, and clean reads were aligned to the reference using HISAT2 (v2.0.5) (Kim et al.2015). Gene read counts were calculated with HTSeq (Anders et al.2015). Differentially expressed genes (DEGs) were identified using the DESeq2 R package (v1.16.1), with FPKM (fragments per kilobase of exon per million mapped fragments) values calculated.

DEGs were defined as genes with an adjusted P-value < 0.05 and a fold change ≥ 2, determined using the false discovery rate (FDR) method. Venn diagrams and heatmaps were generated using VennDetail (Hinder et al.2017) and pheatmap (Hu et al.2021) in R, respectively. Gene ontology (GO) and enrichment analyses were performed using ShinyGO 0.76.3 (Ge et al.2020), leveraging GO data from EnsemblPlants resources (Yates et al.2022).

### 2.5. Nanopore-based 16S sequencing

The isolated nodules and roots underwent a thorough washing with running tap water, followed by surface sterilization in a 3% sodium hypochlorite solution for 3 min, and then rinsed three times with sterilized distilled water. Total DNA extraction was performed using the CTAB method as previously described (Clarke, 2009). For PCR reactions, 5 ng of DNA from each sample was used with 16S primers 27F (AGAGTTTGATCMTGGCTCAG) and 1492R (CGGTTACCTTGTTACGACTT) for amplification of the near full-length bacterial 16S rRNA gene. PCR was carried out with the following thermal cycling conditions: initial denaturation at 95°C for 2 min, followed by 30 cycles of denaturation at 95°C for 30 sec, annealing at 60°C for 30 sec, and extension at 72°C for 2 min. This was succeeded by a final extension at 72°C for 10 min and a hold at 4°C. Amplicons from each sample were ligated to pooled barcoded reads using the 16S Barcoding Kit 24 V14 (SQK-16S114.24, Oxford Nanopore Technologies, Oxford, UK) for library construction. Sequencing was performed on a MinION portable sequencer (Oxford Nanopore Technologies) using a flow cell (R10.4.1).

The MinION™ sequencing data (FASTQ files) were processed using the EPI2ME software from Oxford Nanopore Technologies to generate pass reads in FASTQ format with an average quality score exceeding 9. The EPI2ME Fastq Barcoding workflow (Oxford Nanopore Technologies) was employed for trimming adapter and barcode sequences. Principal coordinate analysis, alpha diversity, and relative abundance of the bacterial community were computed using the R packages phyloseq and vegan. Non-parametric Kruskal–Wallis tests were applied to detect differences in taxa abundance at a significance level of 5% (McMurdie and Holmes 2013).

### 2.6. Real-time qPCR analysis of nitrogen-fixing genes in root nodules

Before RNA extraction from pea nodules, we thoroughly removed the surface contaminants by washing them with PBS buffer. RNA extraction and cDNA synthesis were performed followed by SV Total RNA Isolation System (Promega Corporation, USA) and GoScript™ Reverse Transcriptase (Promega Corporation, USA) kits, respectively. The expressions of *NifA* (nitrogenase Fe protein regulatory protein), *NifD* (nitrogenase FeMo protein alpha subunit), and *NifH* (nitrogenase FeMo protein beta subunit) associated with the nitrogen fixation process in diazotrophic bacteria in nodules were then studied using gene-specific primers (Supplementary Table S2) with the CFX96 Real-Time system (Bio-Rad, USA). The qPCR reaction was conducted under the following conditions: initial denaturation at 95°C for 3 min, followed by 40 cycles consisting of denaturation at 95°C for 5 sec, annealing at 56°C for 30 sec, and extension at 60°C for 5 min. Expression levels of target genes were analyzed using the dd^-ΔCt^ method (Livak and Schmittgen, 2001), with normalization to *PsGAPDH*.

### 2.7. Amplicon sequencing of 16S and ITS microbial community

Amplicon sequencing targeting the 16S (bacteria) and ITS (fungi) gene was conducted to analyze microbial communities in the root samples. The root cleaning protocol involved vortexing the samples in sterile phosphate-buffered saline (PBS) for 10 sec, followed by two rinses with sterile water to eliminate surface contaminants. Approximately 0.2 g of cleaned root tissue was utilized for DNA extraction, adhering to the manufacturer’s instructions provided with the Wizard® Genomic DNA Purification Kit (Promega Corporation, United States). This extraction process included treatments with RNase and Proteinase K to remove potential RNA and protein contaminants, respectively.

The amplicon library for the 16S rRNA gene was prepared using the primer pair 341F (CCTACGGGNGGCWGCAG) and 805R (GACTACHVGGGTATCTAATCC). Sequencing was conducted on the Illumina Novaseq 6000 (PE250). Raw sequencing reads were processed and trimmed using the cutadapt software (Martin, 2011), followed by analysis with the DADA2 pipeline (Callahan et al. 2016). Sequences identified as originating from mitochondria and chloroplasts were excluded from the dataset. Taxonomic assignments for amplicon sequence variants (ASVs) were conducted using the UNITE fungal reference database (Nilsson et al. 2019) for internal transcribed spacer (ITS) regions and version 138 of the Silva Project (Quast et al. 2013) for 16S rRNA sequences. Principal Coordinate Analysis (PCoA) was performed by calculating Bray-Curtis distance matrices based on Hellinger-transformed counts, utilizing the phyloseq and vegan packages in R. Non-parametric Kruskal-Wallis tests were employed to assess differences in taxa abundance. ASVs were classified into various taxonomic levels, and comprehensive data analyses—including principal component analysis, alpha diversity, beta diversity, and relative abundance assessment—were conducted at a significance threshold of P < 0.05 using the phyloseq package in R (McMurdie and Holmes 2013).

### 2.8. Determinations of siderophore levels in the rhizosphere

The siderophore content in rhizosphere soil was quantified using a chrome azurol S (CAS) assay, following the protocol described by Himpsl and Mobley (2019). Briefly, rhizosphere soil samples were homogenized in 80% methanol and centrifuged at 10,000 rpm for 15 minutes. The supernatant (500 μl) was then combined with 500 μl of CAS solution and incubated at room temperature for 5 minutes. Absorbance was measured at 630 nm, with a reference consisting of 1 ml of CAS reagent alone. Siderophore levels were calculated using the formula: % siderophore unit = [(Ar – As) / Ar] x 100, where Ar is the absorbance of the reference (CAS reagent only) and As is the absorbance of the sample (CAS reagent + soil sample).

### 2.9. Colonization efficiency of C. globosum in the roots

The establishment and colonization of *C. globosum* in the roots were evaluated by measuring the relative abundance by DNA-based qPCR using the following primers: forward – AGCAAAGCAAACACTCTTGGCT and reverse – CACGCCATCACTCGGTCGT) as previously described (Nakayama et el. 2013). Root samples were washed twice in sterile phosphate-buffered saline, followed by brief vortexing and two rinses with sterile water. DNA was extracted from approximately 0.2 g of root tissue using the CTAB (cetyl trimethylammonium bromide) method (Clarke, 2009). The extracted DNA was quantified using a NanoDrop ND-1000 Spectrophotometer (Wilmington, USA) and standardized to equal concentrations across all samples. Quantitative PCR (qPCR) was performed using a CFX96 Touch Real-Time PCR Detection System (Bio-Rad, USA), with PsGAPDH serving as the internal control for relative quantification using the 2-ΔΔCT method (Livak and Schmittgen, 2001).

### 2.10. Data analysis

All data were collected from three independent biological replicates for each treatment. The statistical significance of each variable was assessed through Student’s *t-test* and ANOVA followed by Duncan’s Multiple Range Test (DMRT), where appropriate. Graphical presentations were prepared using ggplot2 in R program.

## 3. Results

### 3.1. Measurement of morphological, photosynthesis and nutrient parameters

The cultivation of garden peas exposed to high pH resulted in impaired aboveground growth (Fig. 1A-1B). The garden pea plants showed a significant decline in root length, root dry weight, shoot height and shoot dry weight due to high pH relative to untreated controls (Fig. 1C-F). Further, the chlorophyll score, Fv/Fm, and Pi_ABS exhibited a significant decline in leaves of garden pea under high pH conditions in soil compared to untreated plants (Fig. 1G–1I). Furthermore, ICP-MS analysis showed a significant decrease in Fe, Zn, Mn, and S in both root and shoot of pea plants exposed to high pH in contrast to controls (Table 1). However, Mg and Ca levels exhibited no significant alterations in response to high pH in comparison to controls (Table 1).

**Fig 1.**
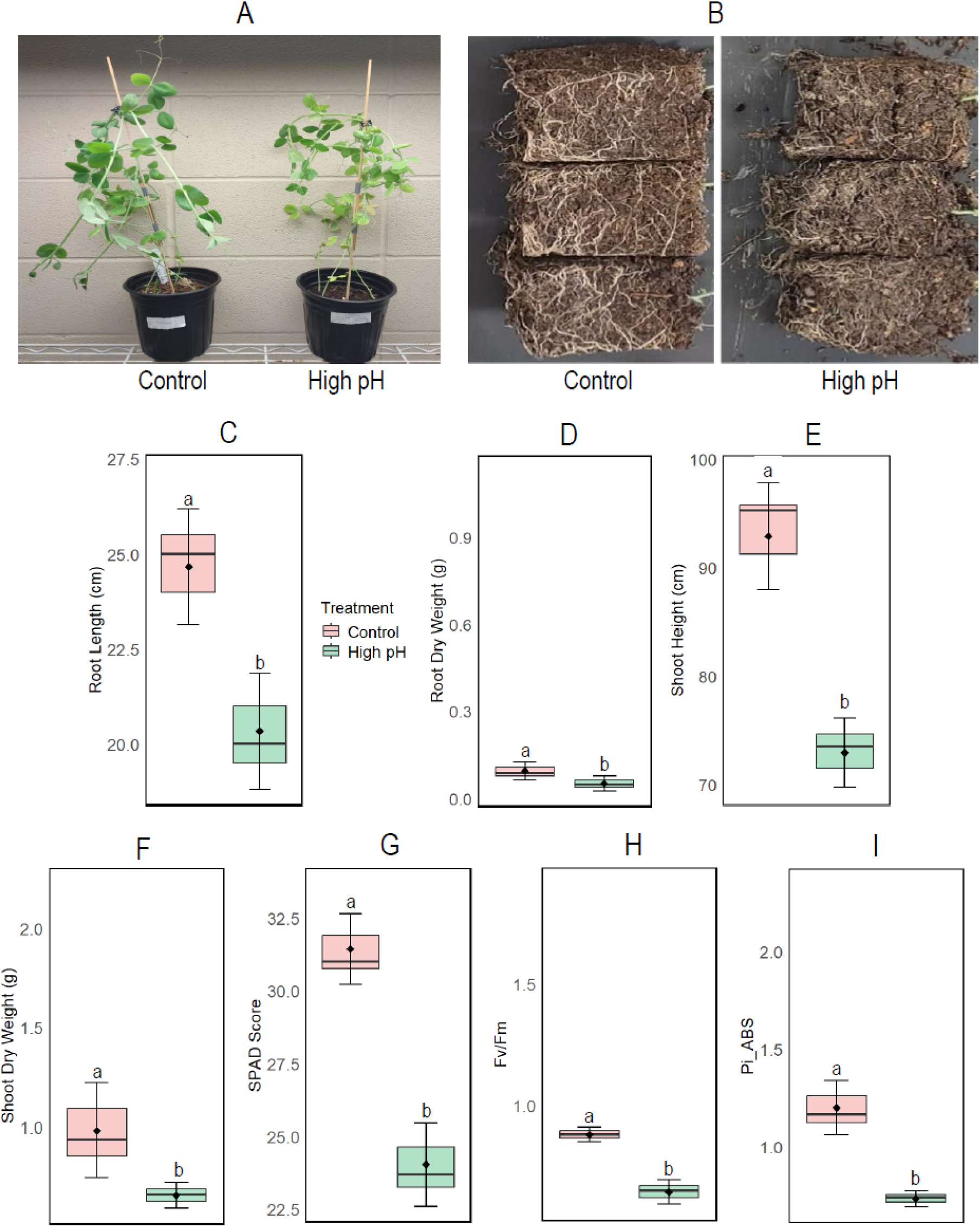
Morphological and physiological responses of garden peas grown in control and high pH conditions: aboveground phenotype. (A), belowground phenotypes (B), root length (C), root dry weight (D), shoot length (E), and shoot dry weight (F), leaf SPAD score (G), leaf Fv/Fm (maximal quantum yield of PS II (H) and leaf Pi_ABS (photosynthesis performance index) (I). Plans were cultivated for 5 weeks in greenhouse conditions. Different letters above the bar indicate significant differences between the control and high pH conditions at a *P* <0.05 level. The data represents the means ± SD of three independent biological samples (*n* = 3).

**Table 1.** ICP-MS analysis of nutrients in the root and shoot of garden peas grown in control and high-pH soils. Different letters indicate significant differences between the control and high pH conditions at a *P* <0.05 level. The data represents the means ± SD of three independent biological samples (*n* = 3).

| Elements | Root |  | Shoot |  |
| --- | --- | --- | --- | --- |
|  | Control | High pH | Control | High pH |
| Fe (ppm) | 384.4±8.7<sup>a</sup> | 287.6±8.0<sup>b</sup> | 92.9±10.3<sup>a</sup> | 58.2±1.2<sup>b</sup> |
| Zn (ppm) | 126.9±15.1<sup>a</sup> | 90.2±5.2<sup>b</sup> | 52.2±5.9<sup>a</sup> | 52.7±4.9<sup>a</sup> |
| Mn (ppm) | 37.7±2.9<sup>a</sup> | 22.4±1.9<sup>b</sup> | 37.1±2.4<sup>a</sup> | 18.3±1.2<sup>b</sup> |
| S (%) | 1.2±0.2<sup>a</sup> | 0.7±0.1<sup>b</sup> | 0.6±0.1<sup>a</sup> | 0.3±0.01<sup>b</sup> |
| Mg (%) | 0.2±0.01<sup>a</sup> | 0.2±0.01<sup>a</sup> | 0.2±0.01<sup>a</sup> | 0.2±0.01<sup>a</sup> |
| Ca (%) | 0.8±0.1<sup>a</sup> | 0.7±0.1<sup>a</sup> | 1.2±0.01<sup>a</sup> | 0.4±0.01<sup>a</sup> |

### 3.2. Changes in nitrogen fixation genes, and bacterial community in nodules

We counted the number of nodules in the root which exhibited a significant decline in nodules in pea roots in response to high pH (Fig. 2A). Our qPCR experiments demonstrated that the expression of *NifA* and *NifD* significantly declined in nodules due to high pH relative to the plants cultivated with neutral pH. However, the *NifH* showed a significant upregulation in nodules under soil alkalinity compared to controls (Fig. 2B). We performed Oxford Nanopore Sequencing to analyze the 16S rRNA gene of bacterial communities in nodules collected from pea plants cultivated under control and high pH conditions. Our 16S analysis showed no significant changes in the richness (Chao 1) and diversity (Shannon index) of bacterial communities in root nodules regardless of pH levels in the soil (Fig. 2C-2D). However, the relative abundance analysis showed a significant in *Rhizobium* genera among the top 10 bacterial communities in nodules due to high pH relative to controls (Fig. 2E). At the species level, *Rhizobium indicum*, *Rhizobium leguminosarum,* and *Rhizobium redzepovicii* were the most dominant Rhizobia group among the top 5 bacterial species in pea nodules. Furthermore, all three species showed a significant reduction in the relative abundance of nodules under high pH conditions in contrast to untreated controls (Fig. 2F).

**Fig 2.**
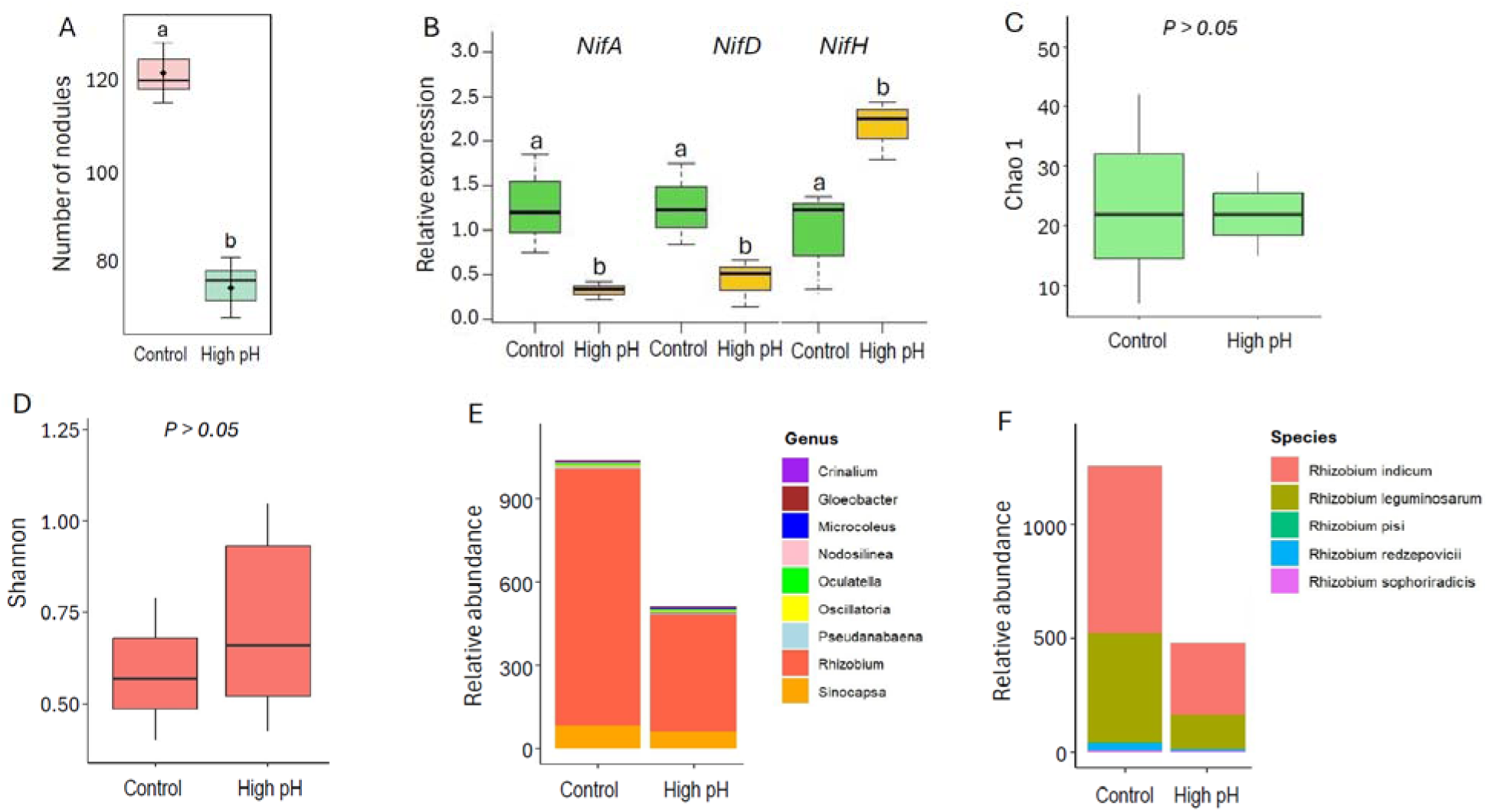
Effect of high pH on the root nodule numbers. (A), relative expression of *nif* genes (B): *nifA, nifD,* and *nifH*, Chao 1 index (C), Shannon-index (D) for bacterial communities of root nodules, relative abundance of top 10 bacterial genus (E) and relative abundance of top 5 Rhizobia species in root nodules (F) of garden peas. Plans were cultivated for 5 weeks in greenhouse conditions. Different letters above the bars indicate significant differences between the control and high pH conditions at a P <0.05 level. The data represents the means ± SD of three independent biological samples (*n* = 3).

### 3.3. RNA-sequencing analysis for differentially expressed genes (DEGs)

We performed RNA-seq analysis to evaluate the differentially expressed genes in the roots of garden pea exposed to high pH. Principal Component 1 (PC1) accounted for 57.4% of the variability, while PC2 explained 27.4% of the variability for differentially expressed genes in garden pea roots grown under control and high pH conditions (Fig. 3A). The RNA-seq analysis unveiled extensive shifts in the transcriptome of garden pea roots under high pH, with 770 differentially expressed genes (DEGs) exhibiting significant upregulation and 842 genes displaying downregulation (Fig. 3B).

**Fig. 3.**
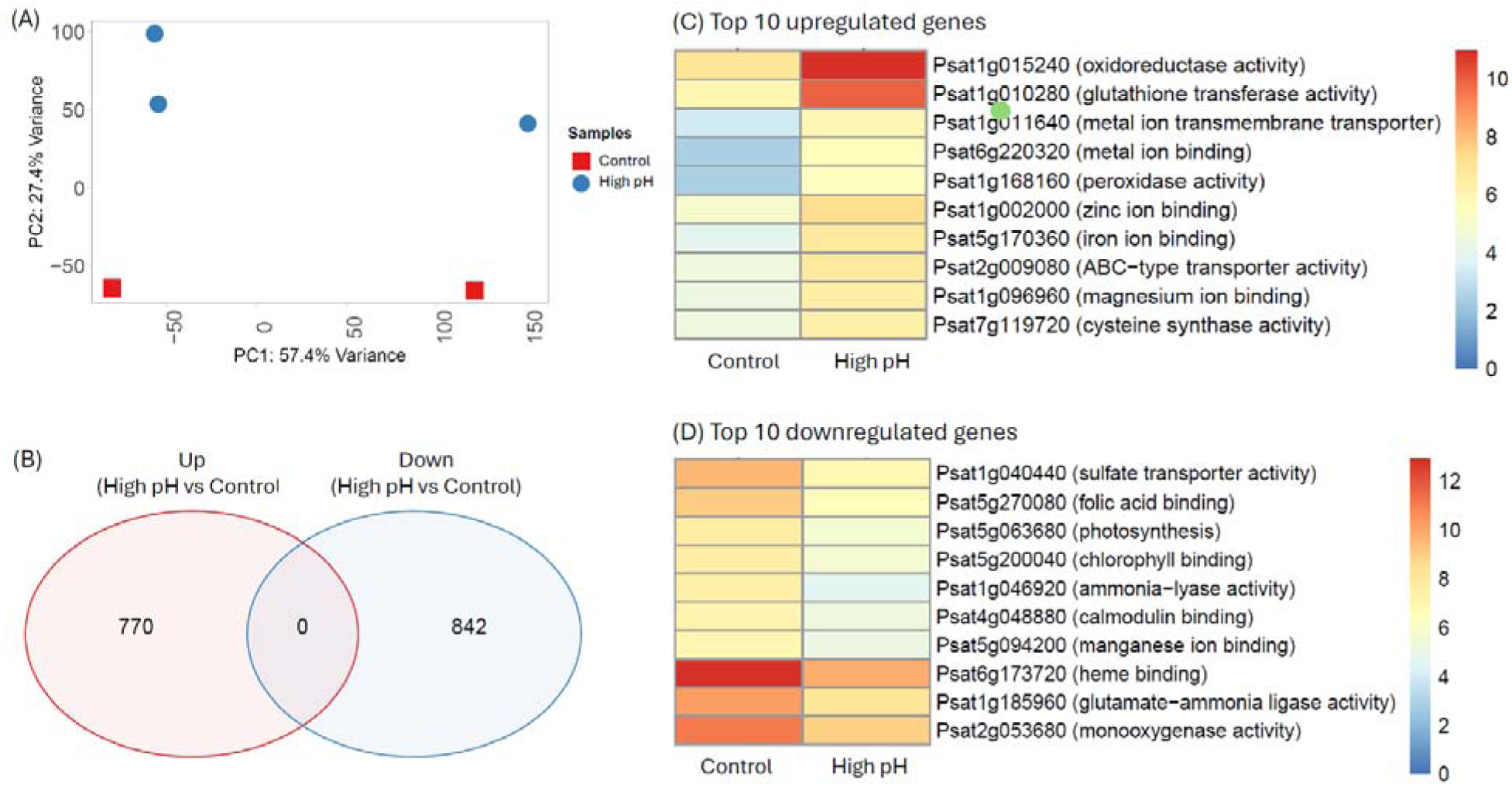
Analysis of RNA-sequencing data for differentially expressed genes (DEGs): (A) principal component analysis of DEGs, (B) vein diagram (>Log2 fold, p<0.05) of upregulated and downregulated genes and Heatmaps for top 10 upregulated genes (A) and for top 10 downregulated genes (C) of garden peas grown in high pH vs. control. Plans were cultivated for 5 weeks in greenhouse conditions.

Subsequent functional annotations of these DEGs were conducted using the EMBL-EBI database specific to garden peas (*Pisum sativum* – Ensembl Genomes 59). This was followed by the generation of heatmaps spotlighting the top 10 upregulated and downregulated (Log2 fold change, p<0.05) genes under high pH conditions compared to the control (Fig. 3C-3D, Supplementary Table S3). Of the upregulated genes, high pH caused a significant upregulation of genes linked to oxidoreductase activity (*Psat1g015240*) and glutathione transferase activity (*Psat1g010280*) in garden peas grown under high pH (Fig 3C). Furthermore, genes associated with diverse functions such as metal ion transmembrane transport (*Psat1g011640*), metal ion binding (*Psat6g220320*), peroxidase activity (*Psat1g168160*), zinc ion binding (*Psat1g002000*), Fe ion binding (*Psat5g170360*), ABC-type transporter activity (*Psat2g009080*), magnesium ion binding (*Psat1g096960*), and cysteine synthase activity (*Psat7g119720*) demonstrated significant upregulation under high pH conditions compared to the control (Fig. 3C). In contrast, genes responsible for sulfate transporter activity (*Psat1g040440*), folic acid binding (*Psat5g270080*), photosynthesis (*Psat5g063680*), chlorophyll-binding (*Psat5g200040*), ammonia-lyase activity (*Psat1g046920*), calmodulin binding (*Psat4g048880*), manganese ion binding (*Psat5g094200)*, heme binding (*Psat6g173720*), glutamate-ammonia ligase activity (*Psat1g185960*), and monooxygenase activity (*Psat2g053680*) were significantly downregulated under high pH conditions compared to the controls (Fig. 3D, Supplementary Table S3).

### 3.4. Gene enrichment analysis

Functional enrichment analysis was conducted on both upregulated and downregulated genes using ShinyGO 0.80 bioinformatics tools. Of the biological processes, the most enriched downregulated pathway was photosynthesis, followed by the metabolic process involving porphyrin-containing compounds (Fig. 4A). Similarly, processes such as tetrapyrrole metabolism, tetrapyrrole biosynthesis, and chlorophyll metabolism were also downregulated under high pH conditions. Conversely, certain biological processes like amide biosynthesis, peptide biosynthesis, translation, cellular amide metabolism, peptide metabolism, and organonitrogen compound metabolism were upregulated in garden peas subjected to high pH compared to the control (Fig. 4A). Further, cellular component metabolic processes related to plastids, chloroplasts, cytoplasm, thylakoids, photosynthetic membranes, photosystems, membrane protein complexes, photosystem I, cellular anatomical entities, and membranes were downregulated, while intracellular non-membrane-bounded organelles, ribosomes, intracellular organelles, organelles, intracellular anatomical structures, cellular anatomical entities, cellular components, nucleosomes, and chromatin were upregulated (Fig. 4B). Of the molecular functions of DEGs, genes associated with structural molecular activity, ribosomal structural constituents, chromatin structural constituents, nucleoside phosphate binding, nucleotide binding, small molecule binding, anion binding, purine nucleotide binding, purine ribonucleoside triphosphate binding, and purine ribonucleotide binding were upregulated in garden peas grown under high pH conditions. Conversely, molecular functions related to oxygen binding, carboxylic acid binding, fatty acid binding, metal ion binding, cation binding, amylase binding, and lipid binding were downregulated (Fig. 4C). According to KEGG pathway analysis, pathways including biosynthesis of secondary metabolites, metabolic pathways, porphyrin metabolism, nitrogen metabolism, alanine, aspartate, and glutamate metabolism, biosynthesis of amino acids, sulfur metabolism, photosynthesis, photosynthetic antenna processes, and the MAPK signaling pathway were downregulated during high pH. Conversely, pathways such as valine, leucine, and isoleucine biosynthesis, oxocarboxylic acid metabolism, biosynthesis of amino acids, and ribosomes were upregulated (Fig. 4D).

**Fig 4.**
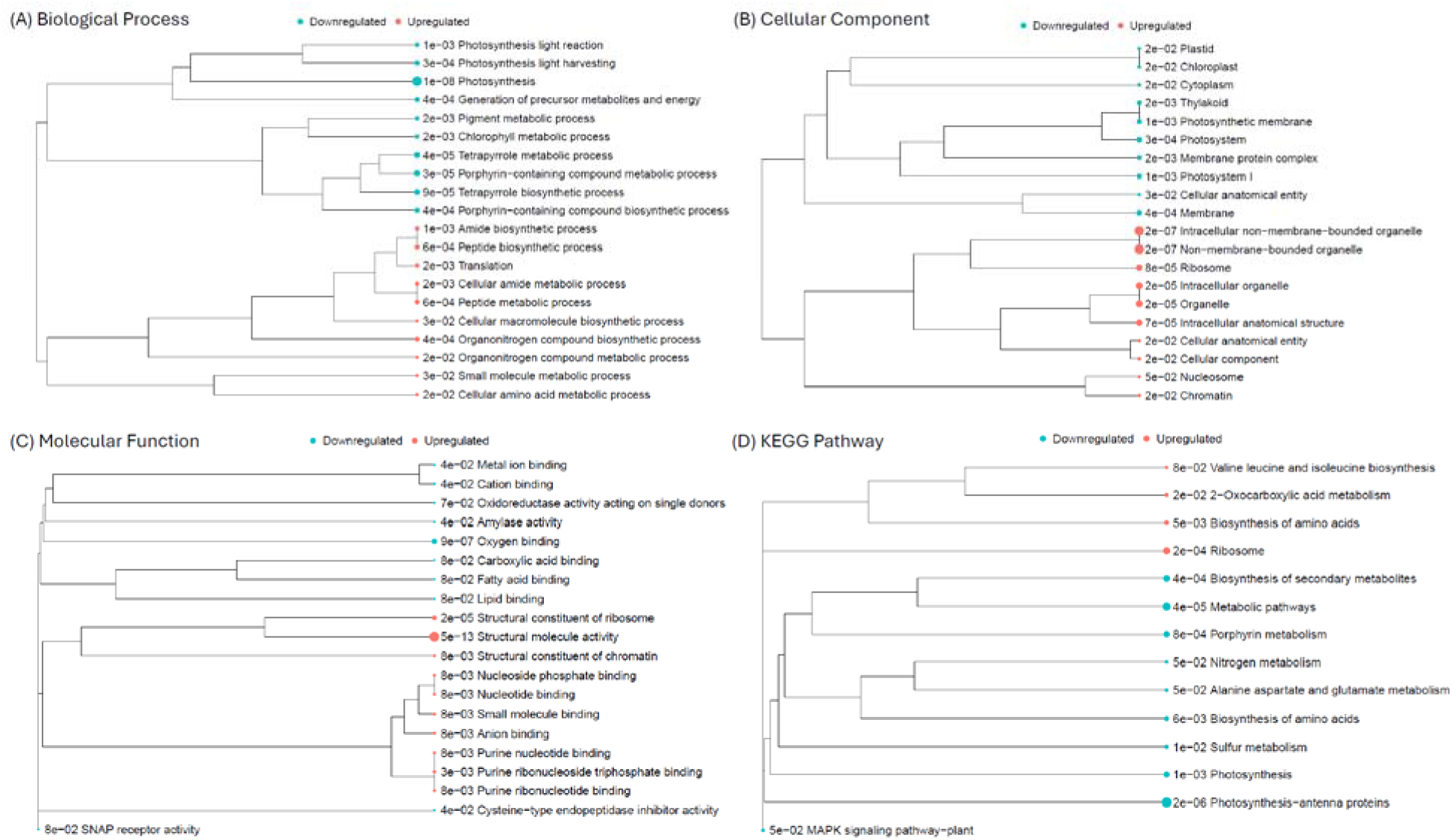
Gene enrichment analysis for the upregulated and downregulated differentially expressed genes (DEGs) of garden peas roots cultivated in high pH and neutral pH conditions: (A) enrichment analysis of upregulated and downregulated genes of roots of garden peas grown in Fe– and control conditions for biological process, (B) enrichment analysis of upregulated and downregulated genes of roots of garden peas grown in high pH and control conditions for cellular components, (C) enrichment analysis of upregulated and downregulated genes of roots of garden peas grown in Fe– and control conditions for molecular functions, and (D) KEGG pathway analysis upregulated and downregulated genes of roots of garden peas grown in high pH and control conditions.

**Fig. 5.**
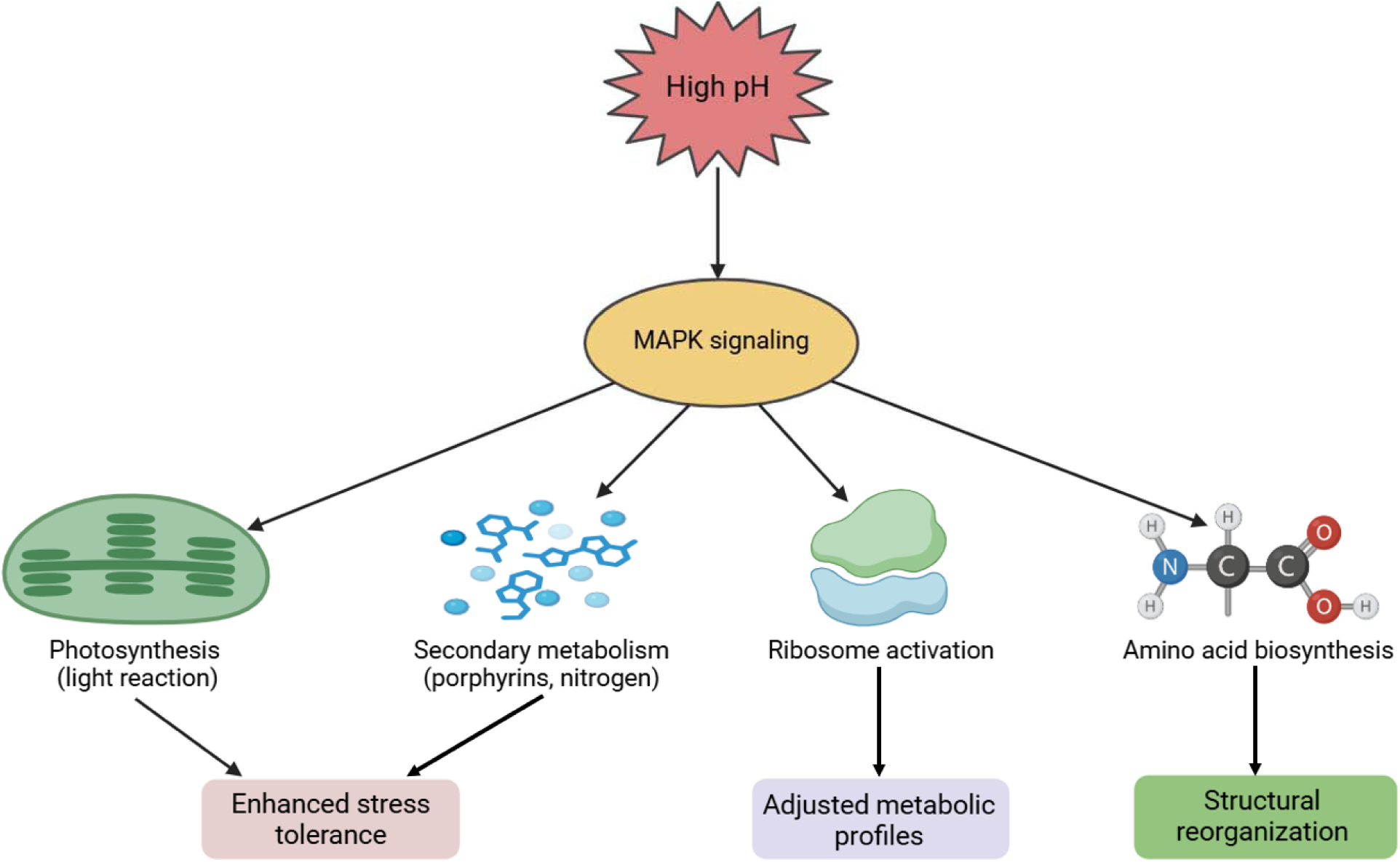
A model summarizing enriched pathways under high pH stress, highlighting the role of MAPK signaling and its associated pathways. Under high pH conditions, MAPK signaling is activated, influencing several key metabolic and physiological pathways. These include enhancement of photosynthesis (light reactions), modulation of secondary metabolism (such as porphyrin and nitrogen-related pathways), ribosome activation to adjust metabolic profiles, and regulation of amino acid biosynthesis, leading to structural reorganization. Collectively, these processes contribute to enhanced stress tolerance and adaptation in garden pea under high pH conditions.

### 3.5. Changes in root microbial community due to high pH

The PCoA represented different dimensions of variation indicating a clear separation between sample groups (Fig. 6A). For bacteria, axis 1 explains 82.9% of the variance, while Axis 2 explains 9.0%, highlighting distinct clustering between the control and high pH treatments. Similarly, for fungi, Axis 1 explains 40.2% of the variance, and Axis 2 explains 36.5%, with a clear separation between control and high pH stress (Fig. 6A). In this study, the Chao1 index (*p* = 0.28) and Simpson index (*p* = 0.11) indicate no significant differences between the control and high pH conditions, suggesting that bacterial richness and evenness are not markedly affected by high pH stress. For fungi, the Chao1 index (*p* = 0.041) shows significantly reduced fungal richness under high pH stress compared to the control. The Simpson index (*p* = 0.045) also reveals a significant reduction in fungal evenness under high pH stress (Fig. 6B). The relative abundance of bacterial orders in the roots differs significantly between control and high pH conditions, with notable changes in specific orders. We found that *Burkholderiales* significantly increased under high pH stress, while *Micrococcales* showed a significant decrease, suggesting sensitivity to pH stress (Fig. 6C). Also, dominant orders such as *Rhizobiales* and *Pseudomonadales* remained relatively stable across treatments, maintaining a consistent proportion in the microbial community (Fig. 6C). In this study, *Variovorax* exhibited as the dominant genus in both conditions, with a significant increase under high pH stress (Fig. 6D). Similarly, *Shinella* showed a significant increase under high pH, while *Pararhizobium* decreased significantly. Other genera, such as *Pseudomonas* and *Massilia*, remained relatively stable across treatments, while minor, non-significant variations are observed in genera like *Streptomyces* and *Flavobacterium* (Fig. 6D). In this study, the fungal community composition at the order level is dominated by *Sordariales* in both control and high pH conditions (Fig. 6E). The relative abundance of the dominant orders, including *Sordariales*, remained consistent across treatments. In addition, minor orders, such as *Hypocreales*, *Pleosporales*, and *Agaricales*, were present in smaller proportions and exhibited slight non-significant shifts between control and high pH conditions, while orders like *Eurotiales* and *Ascosphaerales* remained relatively stable (Fig. 6E). Furthermore, high pH stress induced significant changes in specific genera, notably decreasing the abundance of *Cladosporium*, while *Chaetomium* showed a significant increase under high pH. Further, minor genera such as *Talaromyces*, *Penicillium*, and *Serendipita* exhibited smaller, non-significant shifts in abundance between treatments (Fig. 6F). In addition, we also determined the siderophore levels in the rhizosphere soil which showed a significant increase due to high pH compared to untreated controls (Fig. 6G).

**Fig 6.**
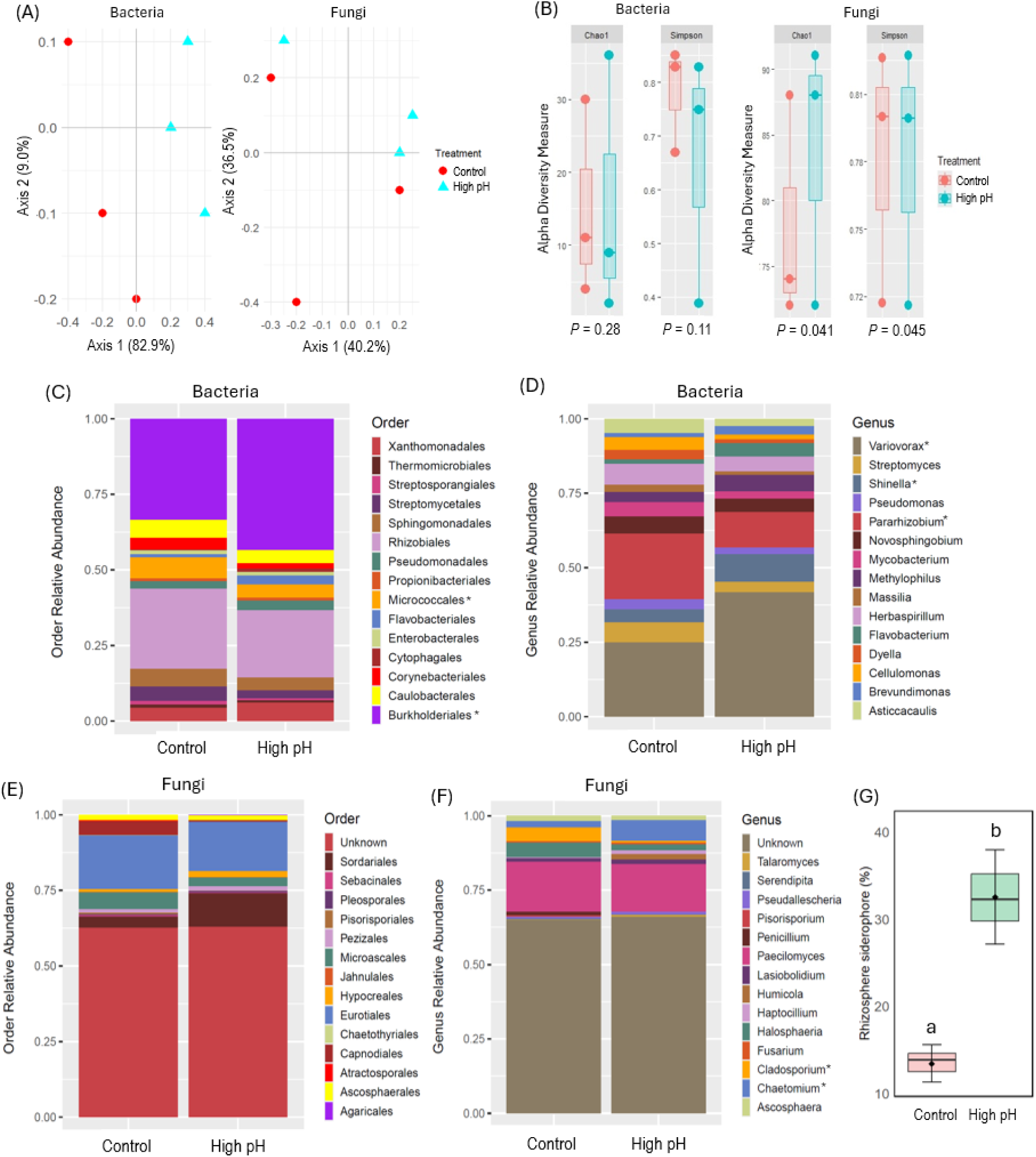
Principal coordinate analysis (PCoA) showing the community compositions assignments of bacterial 16S genes. (A), alpha diversity measures (B): Chao 1, and Simpson index, order-level relative abundance of bacterial taxa (C), genus-level relative abundance of bacterial taxa (D), order-level relative abundance of fungal taxa (E), genus-level relative abundance of fungal taxa in the roots and rhizosphere siderophore (%) of pea plants (G) cultivated in the absence (control) or presence of high pH. Taxa marked with asterisks indicate significant differences between the two growth conditions. The data represents the means ± SD of three independent biological samples (*n* = 3).

### 3.6. Effect of microbial inoculants on Fe-deficient plants

Based on the amplicon sequencing, *C. globosum* and *V. paradoxus* were selected as representative species for the microbial consortia study given their potential functional roles in plant growth promotion. We studied the effects of microbial inoculants consisting of single and combined pure cultures of *C. globosum* and *V. paradoxus* on the growth and physiological parameters of plants under control and high pH conditions. Shoot height and shoot dry weight were significantly reduced under high pH conditions compared to the control. Under high pH stress, inoculation of plants with *C. globosum* showed a significant improvement in these shoot parameters compared to non-inoculated plants (Fig. 7A-7B). However, plants inoculated with *V. paradoxus* and *C. globosum* + *V. paradoxus* under high pH conditions showed no significant changes compared to the plants solely cultivated under high pH. Further, plants inoculated with either *C. globosum* or *V. paradoxus* or in combination showed similar trend in shoot height and shoot dry weight to those of untreated controls (Fig. 7A-7B). High pH stress significantly reduced root length. However, inoculation with *C. globosum*, *V. paradoxus*, or their combination (*C. globosum* + *V. paradoxus*) significantly improved the root length relative to the control (Fig. 7C). Further, plants inoculated with either *C. globosum* or *V. paradoxus* or in combination without high pH stress showed similar root length to those of untreated controls (Fig. 7C). As expected, high pH stress caused a significant decline in root fresh weight (Fig. 7D). However, plants inoculated with either with *C. globosum* or *V. paradoxus* or in combination in response to high pH showed no significant improvement in root fresh weight to those plants cultivated with high pH. Plants inoculated solely with microbial inoculants showed a similar root fresh to that of controls (Fig. 7D). In this study, high pH stress caused a significant reduction in leaf SPAD score compared to controls (Fig. 7E). Plants inoculated with *C. globosum* and *V. paradoxus* either alone or in combination under high pH showed a significant increase in SPAD score relative to the plants solely cultivated with high pH conditions. Plants solely inoculated with *C. globosum* showed a similar SPAD score in the leaves to those of plants stressed with high pH but inoculated with microbial inoculants. However, plants inoculated with *V. paradoxus* and *V. paradoxus* + *V. paradoxus* showed similar SPAD scores to those of controls (Fig. 7E). High pH stress also caused a significant decrease in leaf Fv/Fm compared to controls (Fig. 7F). Plants inoculated with *C. globosum* and *V. paradoxus* alone or in combination with or without high pH showed a significant increase in Fv/Fm compared to the plants solely cultivated under high pH (Fig. 7E).

**Fig. 7.**
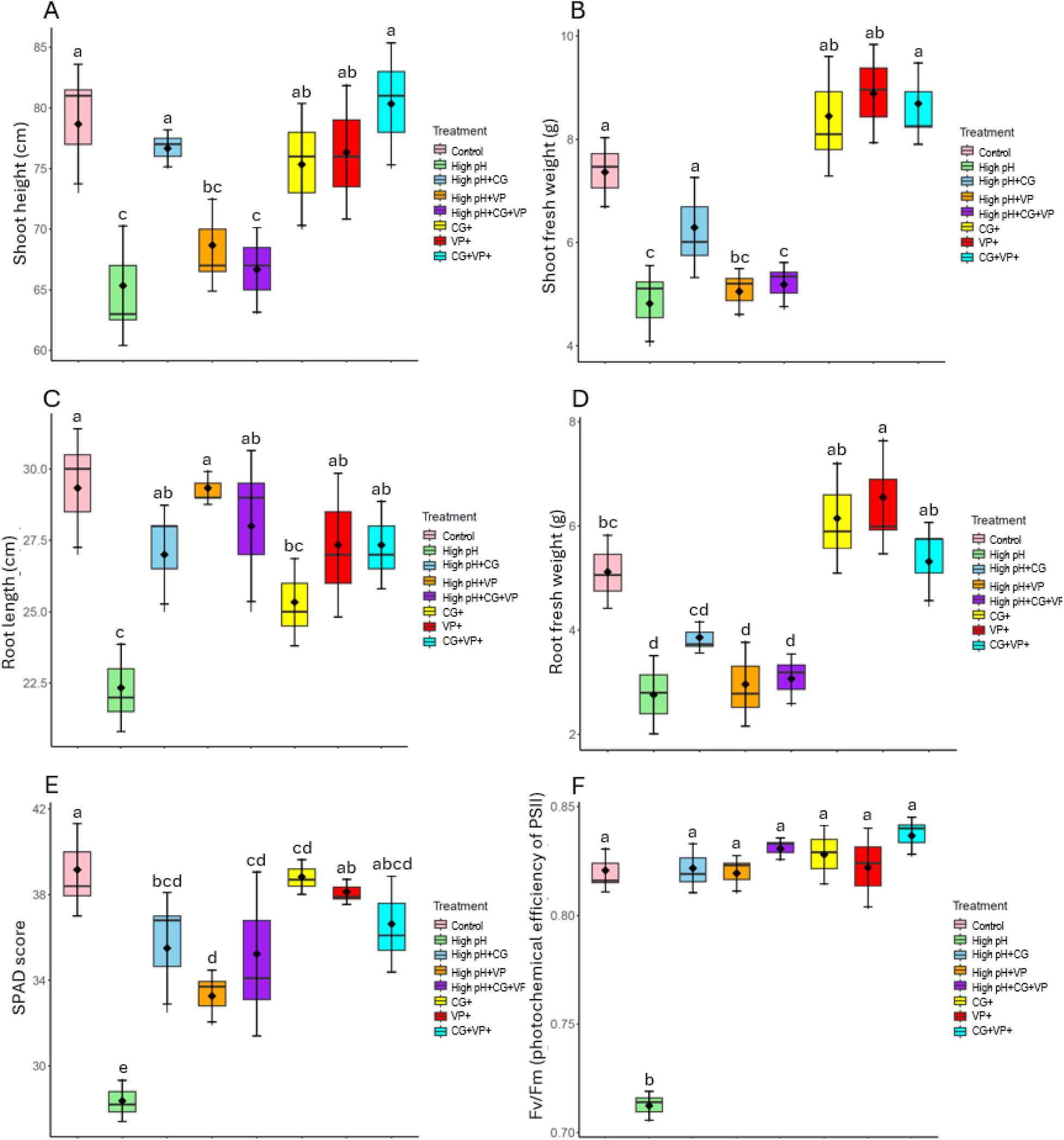
Effects of *Chaetomium globosum* (CG) and *Variovorax paradoxus* (VP) inoculation on morphological and physiological parameters of garden pea plants under control and high pH conditions: (A) shoot height, (B) shoot fresh weight, (C) root length, (D) root fresh weight, (E) SPAD score (chlorophyll content indicator), and (F) Fv/Fm (photochemical efficiency of PSII) Different letters above the bars indicate statistically significant differences among treatments (*p* < 0.05). The data represents the means ± SD of three independent biological samples (*n* = 3).

Based on morphological and physiological parameters, we selected *C. globosum* as a potential inoculant for promoting stress resilience in garden pea exposed to high pH. We determined root colonization efficiency and performed Oxford Nanopore sequencing to investigate whether inoculation with *C. globosum* alters the 16S microbial communities in the roots. In this study, high pH stress caused no significant changes in the relative abundance of *C. globosum* in the root compared to controls (Fig. 8A). The inoculation of plants with *C. globosum* under high pH demonstrated a significant increase in the relative abundance of *C. globosum* relative to high pH stressed plants. Plants inoculated with *C. globosum* without high pH showed a similar relative abundance of *C. globosum* compared to the plants cultivated under control and high pH conditions (Fig. 8A).

**Fig. 8.**
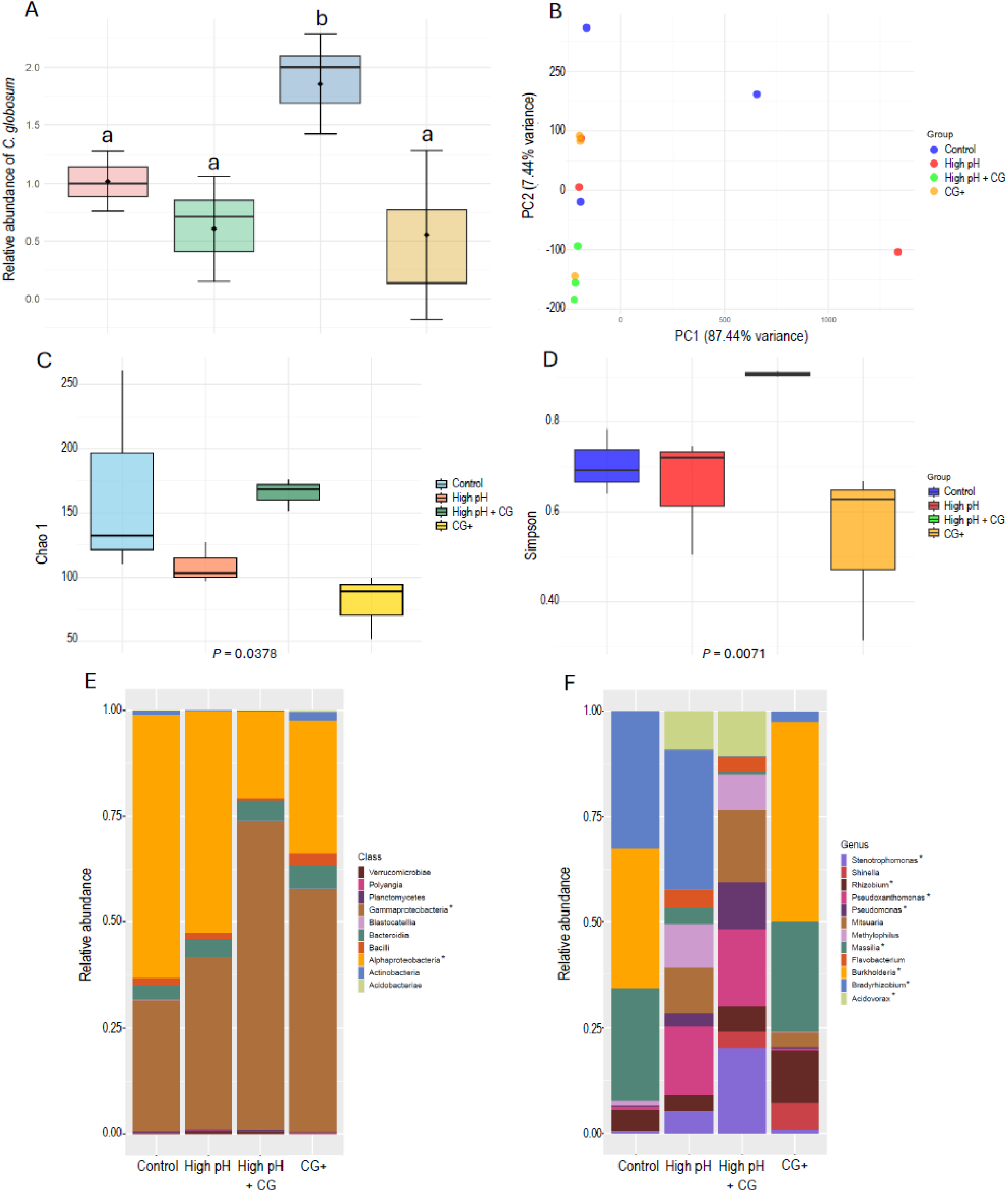
Effects of *Chaetomium globosum* (CG) on microbial community composition and diversity in garden pea roots under high pH stress and control conditions: (A) relative abundance of *C. globosum*, (B) principal component analysis (PCA) of microbial community composition, (C) Chao 1 index indicating microbial richness across treatments, (D) Simpson index reflecting microbial diversity, (E) relative abundance of microbial classes across treatments, (F) relative abundance of microbial genera. Different letters above the bars indicate statistically significant differences among treatments (*p* < 0.05), where appropriate. The data represents the means ± SD of three independent biological samples (*n* = 3).

The PCA demonstrates that high pH stress induces a significant shift in microbial community composition (separation along PC1), while the application of *C. globosum* partially mitigates the stress (Fig. 8B). It also showed the distinct clustering of the control and *C. globosum* + treatments indicating that microbial community profiles are strongly influenced by both stress and *C. globosum* inoculation (Fig. 8B). The Chao 1 index, which measures microbial richness, reveals that high pH stress significantly reduces microbial richness compared to the control. However, the introduction of *C. globosum* under high pH stress substantially restores microbial richness indicating that *C. globosum* plays a critical role in mitigating the adverse effects of high pH stress on microbial diversity (Fig. 8C). Further, *C. globosum* applied to high pH stress maintains microbial diversity at a level comparable to the control as evident by the Shannon index (Fig. 8D). The relative abundance of microbial classes in the roots varied significantly across treatments, with high pH stress altering the community composition by reducing key classes such as *Alphaproteobacteria*. The inoculation of *C. globosum* under high pH substantially increased the relative abundance of *Gammaproteobacteria* while reducing the *Alphaproteobacteria* (Fig. 8E). Under non-stress conditions, *C. globosum* selectively promoted certain classes like *Actinobacteria*, further influencing the microbial community structure (Fig. 8E). In addition, high pH stress significantly alters the bacterial composition by reducing the relative abundance of *Rhizobium, Burkholderia* and *Massilia*, while increasing genera like *Acidovorax, Stenotrophomonas* and *Pseudoxanthomonas* (Fig. 8F). Inoculation with *C. globosum* under high pH conditions restored the abundance of *Rhizobium* and further increased the abundance of *Stenotrophomonas*, *Pseudoxanthomonas*, and *Pseudomonas* in the roots relative to the roots of plants exposed to high pH conditions. Under non-stress conditions, *C. globosum* restored the relative abundance of *Massilia* and *Burkholderia* to control levels (Fig. 8F).

## 4. Discussion

High soil pH presents a significant challenge to agricultural productivity, particularly in legume species like garden pea (Goyal et al. 2021). This study provides novel insights into the transcriptomic profile and microbial dynamics associated with high soil pH, highlighting shifts in key nitrogen-fixing bacteria. Furthermore, the observed shifts in root-associated bacterial communities, characterized by the enrichment of genera underscore the potential of these microbes in coping with micronutrient imbalances under high pH conditions. Furthermore, inoculation studies using microbial consortia, including *C. globosum* and *V. paradoxus*, demonstrated their effectiveness in reshaping the root microbiome and enhancing plant resilience under high pH conditions. These findings uncover extensive changes in both the plant’s transcriptional landscape and its microbial associations, offering valuable insights into the mechanisms underlying plant adaptation to high pH stress.

### 4.1. Morphological and photosynthetic impairment

The results of our study provide a comprehensive insight into the physiological and molecular responses of garden peas to high pH stress. Plants grown under high pH conditions exhibited stunted growth compared to their control counterparts. This is consistent with previous studies indicating that iron is crucial for various physiological processes, including chlorophyll synthesis and electron transport in photosynthesis (Aung et al. 2022). We also tested the chlorophyll scoring and OJIP measurements which offered valuable insights into the photosynthetic efficiency and stress responses of the plants (Küpper et al. 2019). The decrease in chlorophyll synthesis and chlorophyll fluorescence parameters underscores the detrimental impact of nutrient imbalance due to high pH on the photosynthetic machinery, leading to reduced energy capture and assimilation, which ultimately manifests as stunted growth and reduced biomass. The significantly lower Fv/Fm ratio highlights the detrimental impact of high pH on the maximal photochemical efficiency of PSII, corroborating findings that micronutrients are essential for optimal photosynthetic performance (Briat et al. 2015). Additionally, the decrease in Pi_ABS in plants exposed to high pH indicates further impairment of the photosynthetic apparatus, reinforcing the critical role of micronutrients in maintaining photosynthetic efficiency (Ravet and Pilon 2013).

### 4.2. Rhizobia dynamics in root nodules in response to high pH

We counted the nodules in pea roots that provided important data on the symbiotic relationship between garden peas and nitrogen-fixing bacteria under high pH conditions. Our results showed a significant reduction in nodule number in pea plants exposed to high pH. This finding aligns with earlier reports suggesting that the imbalance of iron due to high pH in soil, which is vital for the functioning of nitrogenase, the enzyme complex responsible for nitrogen fixation in nodules (Schwember et al. 2019; Brear et al. 2013). The reduced nodule formation could thus be a direct consequence of impaired nitrogen fixation capability by Fe deprivation caused by high pH. Furthermore, the downregulation of *NifA* and *NifD* genes in nodules under high pH conditions supports the idea that iron is essential for nitrogenase function, with the alkalinity-induced micronutrient imbalance leading to reduced enzyme activity. *NifA* is a transcriptional activator that regulates the expression of other *Nif* genes, including *NifD*, under optimal conditions (Zhang et al. 2023). Thus, lower *NifA* expression under high pH likely leads to a cascade effect, reducing the overall expression of the nitrogenase complex components, and thereby impairing nitrogen fixation. Interestingly, the upregulation of *NifH* under high pH could indicate a compensatory response aimed at maintaining nitrogenase activity despite the overall stress condition and its symbionts’ attempt to adjust the nitrogen fixation machinery. This differential gene expression pattern highlights the complex regulatory mechanisms plants and symbiotic bacteria employ to adapt to high pH.

Our 16S rRNA gene analysis revealed no significant changes in the richness and diversity of bacterial communities in root nodules between neutral and high pH conditions. This stability in bacterial community diversity suggests that the overall microbial ecosystem within the nodules remains relatively resilient to alkalinity stress. However, the significant reduction in the relative abundance of *Rhizobium* genera and specific species like *Rhizobium indicum*, *Rhizobium leguminosarum*, and *Rhizobium redzepovicii* under high pH conditions highlights a selective impact of micronutrient availability on key nitrogen-fixing bacteria. These *Rhizobium* species are integral to effective nitrogen fixation, and their reduced abundance could directly contribute to the observed decline in nodule formation and nitrogenase gene expression under high pH in garden pea. Similarly, high pH adversely affected the symbiotic efficiency and nitrogen fixation capacity of *Bradyrhizobium japonicum* and *Mesorhizobium cicero* in legume crops (Merry et al. 2022; Gopalakrishnan et al. 2015). These studies highlight the critical importance of pH for the health and functionality of rhizobia populations in pea nodules. Also, our findings underscore the need for strategies to improve micronutrient availability in soils, which could enhance nitrogen fixation and overall plant productivity in pea crops.

### 4.3. Transcriptional changes in roots

Our RNA-seq analysis has provided a detailed overview of the transcriptional responses of garden pea roots to high pH, revealing significant shifts in gene expression profiles. The principal component analysis indicates that the majority of the variability in gene expression can be attributed to nutrient availability which underscores the profound impact of high pH on the root transcriptome. The identification of 770 upregulated and 842 downregulated genes highlights the extensive transcriptional reorganization occurring in response to high pH. This also suggests a broad reprogramming of metabolic and physiological pathways in response to high pH stress.

The upregulation of genes associated with oxidoreductase activity (*Psat1g015240*) and glutathione transferase activity (*Psat1g010280*) suggests an enhanced oxidative stress response. These genes likely play crucial roles in mitigating the oxidative damage caused by high pH, which is known to disrupt cellular redox homeostasis (Shee et al. 2022; Tewari et al. 2013). Also, the expression and activity of certain oxidoreductases are often upregulated in plants under high pH (Vigani and Murgia 2018). For instance, increased activity of enzymes like ferric-chelate reductase is observed in the roots of peas exposed to lime-induced high pH soil, which aids in the reduction of Fe^3+^ to the more soluble Fe^2+^ form, facilitating its uptake (Kabir et al. 2012). Additionally, other oxidoreductases involved in the ascorbate-glutathione cycle, such as ascorbate peroxidase and glutathione reductase, are crucial for detoxifying ROS produced under Fe stress (Hasanuzzaman et al. 2019). Furthermore, the significant upregulation of genes involved in metal ion transmembrane transport (*Psat1g011640*) and binding (*Psat6g220320*, *Psat1g002000*, *Psat5g170360*) indicates a compensatory mechanism to enhance nutrient uptake and mobilization within the plant. This is consistent with previous findings that plants increase the expression of metal transporter genes to cope with alkalinity-induced iron scarcity (Kobayashi and Nishizawa, 2012). The upregulation of ABC-type transporter activity (*Psat2g009080*) and magnesium ion binding (*Psat1g096960*) also points to broader alterations in nutrient transport and homeostasis under high pH conditions. Conversely, the downregulation of genes associated with sulfate transporter activity (*Psat1g040440*), manganese ion binding (*Psat5g094200*), and ammonia-lyase activity (*Psat1g046920*), suggests an impairment of metabolic processes as well as nutrient imbalance and secondary metabolites in peas in response to high pH. The suppression of mineral uptake genes is consistent with the decreased minerals, as iron and manganese are critical components of chlorophyll and the photosynthetic electron transport chain (Kroh and Pilon 2020; Terry and Low 1982). However, roots are non-photosynthetic organs, so the biological significance of this downregulation may not directly reflect the expression levels of these genes in photosynthetic organs such as leaves. Furthermore, ammonia-lyase enzymes, such as phenylalanine ammonia-lyase (PAL), play critical roles in nitrogen assimilation and the biosynthesis of secondary metabolites, including phenolic compounds (Cass et al. 2015; Kim et al. 2014). Under high pH conditions, the reduced expression of ammonia-lyase genes may indicate that the plant is experiencing a stress-induced imbalance in nitrogen availability, which is consistent with the reduced nodulation and decreased expression of *NifA* and *NifD* genes. High soil pH can limit the solubility and bioavailability of essential nutrients further impairing nitrogen metabolism (Ferrarezi et al. 2022).

Our comprehensive RNA-seq and functional enrichment analysis reveals the intricate molecular and physiological responses of garden pea roots to high pH. The significant differential gene expression, coupled with the detailed functional enrichment analysis, highlights the adaptive strategies employed by plants under nutrient stress. In this analysis, the most enriched downregulated pathway was photosynthesis, which aligns with previous findings indicating that Fe is crucial for chlorophyll synthesis and the photosynthetic electron transport chain (Kroh, & Pilon, 2020). Furthermore, the suppression of tetrapyrrole metabolism, tetrapyrrole biosynthesis, and chlorophyll metabolism pathways suggests a direct consequence of Fe scarcity on the photosynthetic machinery, potentially leading to reduced photosynthetic efficiency and overall plant growth. Conversely, upregulated processes such as amide biosynthesis, peptide biosynthesis, translation, and various metabolic processes indicate a shift towards maintaining essential cellular functions and protein synthesis under stress conditions. This shift could represent a strategic allocation of resources to sustain fundamental metabolic activities and ensure the survival of garden pea exposed to high pH conditions.

The enrichment analysis of cellular component metabolic processes revealed that plastids, chloroplasts, thylakoids, and related components were significantly downregulated, which is consistent with the reduced photosynthetic activity observed under high pH. The downregulation of photosystems and membrane protein complexes highlights the detrimental effects of abiotic stresses on the photosynthetic apparatus (Nouri et al. 2015). In contrast, the upregulation of intracellular non-membrane-bounded organelles, ribosomes, nucleosomes, and chromatin suggests an increased focus on maintaining ribosomal function and chromatin structure. This may indicate an adaptive response to support protein synthesis and genomic integrity despite the challenging conditions posed by high pH. The upregulation of nucleoside phosphate binding and nucleotide binding activities suggests enhanced nucleic acid metabolism and energy transfer processes, which are critical for maintaining cellular function under stress (Witte and Herde 2020). Conversely, the downregulation of functions related to oxygen binding, carboxylic acid binding, fatty acid binding, and metal ion binding reflects the broader impact of high pH on metabolic and binding activities. The reduced expression of genes involved in these functions could contribute to compromised metabolic processes and stress responses.

KEGG pathway analysis further corroborates the significant impact of high pH on metabolic and biosynthetic pathways. Downregulated pathways include biosynthesis of secondary metabolites, metabolic pathways, porphyrin metabolism, and photosynthesis-related pathways, underscoring the metabolic constraints imposed by soil alkalinity. The downregulation of the MAPK signaling pathway suggests potential disruptions in signaling mechanisms critical for stress response and adaptation. The MAPK) signaling pathway serves as a significant route through which plants react to abiotic stressors (Sinha et al. 2011). In contrast, upregulated pathways such as valine, leucine, and isoleucine biosynthesis, oxocarboxylic acid metabolism, and ribosomal pathways indicate a strategic focus on amino acid biosynthesis and protein synthesis. These adaptations may help mitigate the adverse effects of high pH by ensuring the continued production of essential proteins and metabolic intermediates in garden pea. We hypothesis that MAPK signaling mediates adaptive responses in pea roots under high pH by enhancing pathways related to photosynthesis, secondary metabolism, ribosome activation, and amino acid biosynthesis. These processes collectively contribute to enhanced stress tolerance, metabolic adjustments, and structural reorganization, helping plants cope with nutrient limitations (Fig. 5).

### 4.4. The shift in the microbial community in the roots under high pH

While transcriptional changes highlight the plant’s intrinsic mechanisms for coping with high pH stress, the observed shifts in root-associated microbial communities suggest a complementary role of microbes in enhancing stress resilience. Our amplicon sequencing results indicate that high pH stress significantly alters the composition of both bacterial and fungal communities in the roots. Specifically, microbiome analysis suggests no changes in the richness and diversity of bacterial communities in the roots due to high pH. While fungal richness remained unaffected by high pH, a significant reduction in fungal diversity was observed. In general, greater microbial diversity is linked to improved plant fitness and resilience to stress (De Vries et al. 2018). The increase in *Burkholderiales* under high pH to a reorganization of the root microbial community, likely driven by the selective pressures of high pH. *Burkholderiales* are often associated with nutrient cycling and stress tolerance (Liu et al. 2021; Enagbonma et al. 2003). Studies demonstrates that inoculation with *Burkholderia phytofirmans* PsJN enhances salt-stress tolerance in *Arabidopsis thaliana* by priming faster molecular and biochemical responses, including proline accumulation, abscisic acid signaling, ROS scavenging, and ion homeostasis (Pinedo et al. 2015). Given that high pH also disrupted the mineral status in the garden, it is possible that *Burkholderiales* may play a role in mobilizing nutrients to help pea plants cope with high pH-induced stress.

At the genus level, the significant increases in *Variovorax* and *Shinella* suggest that these taxa may have specialized functions in response to high pH, potentially contributing to nutrient acquisition or stress tolerance pathways. Studies showed that *Variovorax boronicumulans* have the potential to help host plants regulate their auxin levels (Sun et al. 2018). *Shinella* can also produce plant growth-promoting substances, such as auxins, which stimulate root development and improve nutrient uptake (Saeed et al. 2021). Auxin facilitates micronutrient uptake specifically Fe in plants by promoting the expression of metal transporter genes and enhancing the growth of root hairs, which increases the plant’s ability to absorb micronutrients from the soil (Zhang et al. 2019; Garnica et al. 2018). While the mechanisms are unclear, another report showed that applying *Variovorax* sp. P1R9, either alone or in plant growth-promoting consortia enhanced salt stress tolerance in wheat (Acuña et al. 2024). Thus, the increased abundance of *Variovorax* and *Shinella* observed in this study suggests its potential role in response to high pH supported by the significant increase in siderophore levels in the rhizosphere. Siderophores are high-affinity Fe-chelating compounds secreted by microbes to sequester Fe from the environment, particularly under Fe scarcity (Kabir and Bennetzen 2024; Bhattacharyya and Jha, 2013). Particularly, *V. paradoxus* is a versatile plant-associated bacterium known for its role in promoting plant growth and enhancing stress tolerance (Sun et al. 2018; Jiang et al. 2012). Since high pH conditions can induce Fe deficiency in plants, as evidenced by our findings, it is plausible that *Variovorax* and *Shinella* may contribute to mitigating this deficiency. In contrast, the decrease in *Rhizobiales*, a group known for its role in nitrogen fixation, may reflect a disruption in plant-microbe symbioses critical for nutrient acquisition under normal conditions. This is consistent with the fact that high pH stress can lead to a disruption in the nodulation process in legumes (Lindström and Mousavi 2020). In this study, certain plant growth-promoting genera, such as *Pseudomonas*, *Novosphinogobium*, and *Herbaspirillum*, exhibited no significant changes in abundance. This suggests that these taxa are either insensitive to nutrient availability disrupted due to soil alkalinity or possess adaptive mechanisms that enable their persistence regardless of the pH levels in the soil. These stable bacterial populations may serve as key functional components of the core microbiome, maintaining critical processes such as nutrient cycling or pathogen suppression irrespective of pH status. *Pseudomonas* bacteria are often considered a core microbiome component for plant growth due to their widespread presence in the root environment (Zhou et al. 2024; Zhuang et al. 2021).

For fungi, *Cladosporium* and *Chaetomium* appear to be significantly affected in the roots of garden pea under high pH treatment. The beneficial effects of *Cladosporium* on plants include the production of secondary metabolites that promote plant growth (Răut et al. 2021; Hamayun et al. 2010). The volatiles released by *Cladosporium cladosporioides* CL-1 have been shown to significantly enhance tobacco growth (Diby and Park, 2013). However, it can also act as a plant pathogen under certain conditions (Rivas and Thomas 2005). In addition, *Chaetomium* is well-documented for producing cellulolytic enzymes and antifungal secondary metabolites (Ibrahim et al. 2021; Shanthiyaa et al. 2014). Studies showed that *C. globosum* enhances the phenylpropanoid pathway in chicory roots, increasing phenylalanine and chicoric acid levels, which are crucial for plant fitness and defense mechanisms (Spinelli et al. 2022). In another study, *C. globosum* was shown to effectively control late wilt disease in maize while also enhancing yield. These findings imply that *Chaetomium* may play critical roles in modulating fungal community dynamics and aiding adaptation in pea plants to high pH environments. Collectively, these findings underscore the dynamic nature of the root microbiome in response to high pH and highlight key microbial taxa that may contribute to plant adaptation under high pH conditions.

### 4.5. Effect of enriched microbiome on pea plants exposed to high pH

In this context, we further investigated the effects of *C. globosum* and *V. paradoxus*, either individually or as combined inoculants, on pea plants. In this study, the strategy of using commercially representative strains of microbes may not accurately represent the original amplicon sequencing data and account for the inherent phenotypic variability among microbial strains. However, this study highlights the potential of microbial inoculants, particularly *C. globosum*, in enhancing stress resilience in garden pea plants exposed to high pH conditions. While all tested inoculants (*C. globosum*, *V. paradoxus*, and their combination) contributed to some degree of stress mitigation, *C. globosum* consistently demonstrated superior efficacy across multiple morphological and physiological parameters. The inability to maintain the persistence of *V. paradoxus* may be due to the use of microbial strains as representatives, which may not accurately reflect the diversity captured in the original amplicon sequencing data (Kabir et al. 2024). Furthermore, the growth promotion efficiency of an individual microbial inoculant may be compromised due to the complex interactions with other microbial species in the soil.

At the microbial level, *C. globosum* uniquely altered root bacterial communities, restoring microbial richness and diversity under high pH stress to levels comparable to control conditions. We found that the inoculation of *C. globosum* helps stabilize bacterial community profiles in the roots of garden pea under high pH stress. *C. globosum* inoculation substantially restored richness, demonstrating its crucial role in counteracting the adverse effects of high pH stress on microbial diversity. Furthermore, the restoration of the Simpson index under high pH stress conditions underscores the importance of microbial inoculation in maintaining a balanced endosphere microbiome in garden pea. We also found that *C. globosum* selectively promoted beneficial microbial classes, such as *Gammaproteobacteria*, and restored the relative abundance of key genera, including *Rhizobium*. *Rhizobium* bacteria play a key role in mitigating mineral stress in plants by increasing the availability of nitrogen and siderophores, which are particularly beneficial in nutrient-limited soil environments (Li et al. 2023; Chieb and Gachomo 2023). As a result, the ability of *C. globosum* to reshape and stabilize the microbial community structure highlights its potential that enhances not only plant performance but also root-associated microbial dynamics under high pH stress in garden pea. Interestingly, we also found the enrichment of *Stenotrophomonas*, and *Pseudomonas* in the roots of pea exposed to high pH, highlighting their role in shaping the microbial community under stress conditions. These genera are known to produce bioactive compounds, and nutrient solubilization (Thounaojam et al. 2018; Chang et al. 2005; Dasila et al. 2023). Several species of *Pseudomonas* (Bais et al. 2006; Zakry et al. 2012) and *Stenotrophomonas* (Islam et al. 2015), have been recognized as plant growth-promoter through various direct and indirect mechanisms which include phosphate solubilization, biological nitrogen fixation, and the production of plant growth regulators and siderophores. Taken together, these findings highlight the ability of *C. globosum* in association with *Stenotrophomonas*, and *Pseudomonas* to mitigate stress-related disruptions in microbial diversity and enhance beneficial taxa, thereby promoting stress resilience in pea plants under high pH conditions. The identification of enriched and helper microbes offers opportunities to develop tailored bioinoculants, paving the way for sustainable crop management in high pH soil. Future studies should focus on the functional characterization of these taxa, particularly in the formulation of microbial inoculants or biofertilizers aimed at enhancing plant tolerance to high pH in garden pea and other legume crops.

## 5. Conclusion

Our study sheds light on the multifaceted responses of garden peas to high pH stress, encompassing physiological, molecular, and microbial dynamics. Plants exposed to high pH exhibited stunted growth, impaired photosynthesis, and decreased chlorophyll synthesis, highlighting the critical role of pH level in soil in various physiological processes. Transcriptomic analysis highlighted the plant’s adaptive responses, with upregulation of oxidative stress-related genes and MAPK signaling pathways, alongside downregulation of nutrient transport and metabolic pathways, reflecting the plant’s attempt to mitigate stress. In root nodules, shifts in nitrogen-fixing bacteria and altered expression of nitrogen-fixation genes demonstrated the impact of high pH on legume-microbe symbiosis. Root microbiome analysis identified enrichment of stress-associated microbes such as *Variovorax*, *Shinella*, and *Chaetomium*, while resilient genera like *Pseudomonas* and *Herbaspirillum* emerged as potential core components for adapting high pH conditions (Fig. 9). Inoculation with *Chaetomium globosum* enhanced growth and increased the abundance of beneficial microbes like *Stenotrophomonas* and *Pseudomonas*, underscoring its collective role in stress mitigation. These findings provide valuable insights into the molecular and microbial mechanisms underlying plant stress responses and highlight opportunities for microbiome-aided strategies. Future research should focus on developing tailored microbial consortia, breeding stress-resilient pea varieties, and field-testing these solutions to optimize legume production in alkaline soils.

**Fig. 9.**
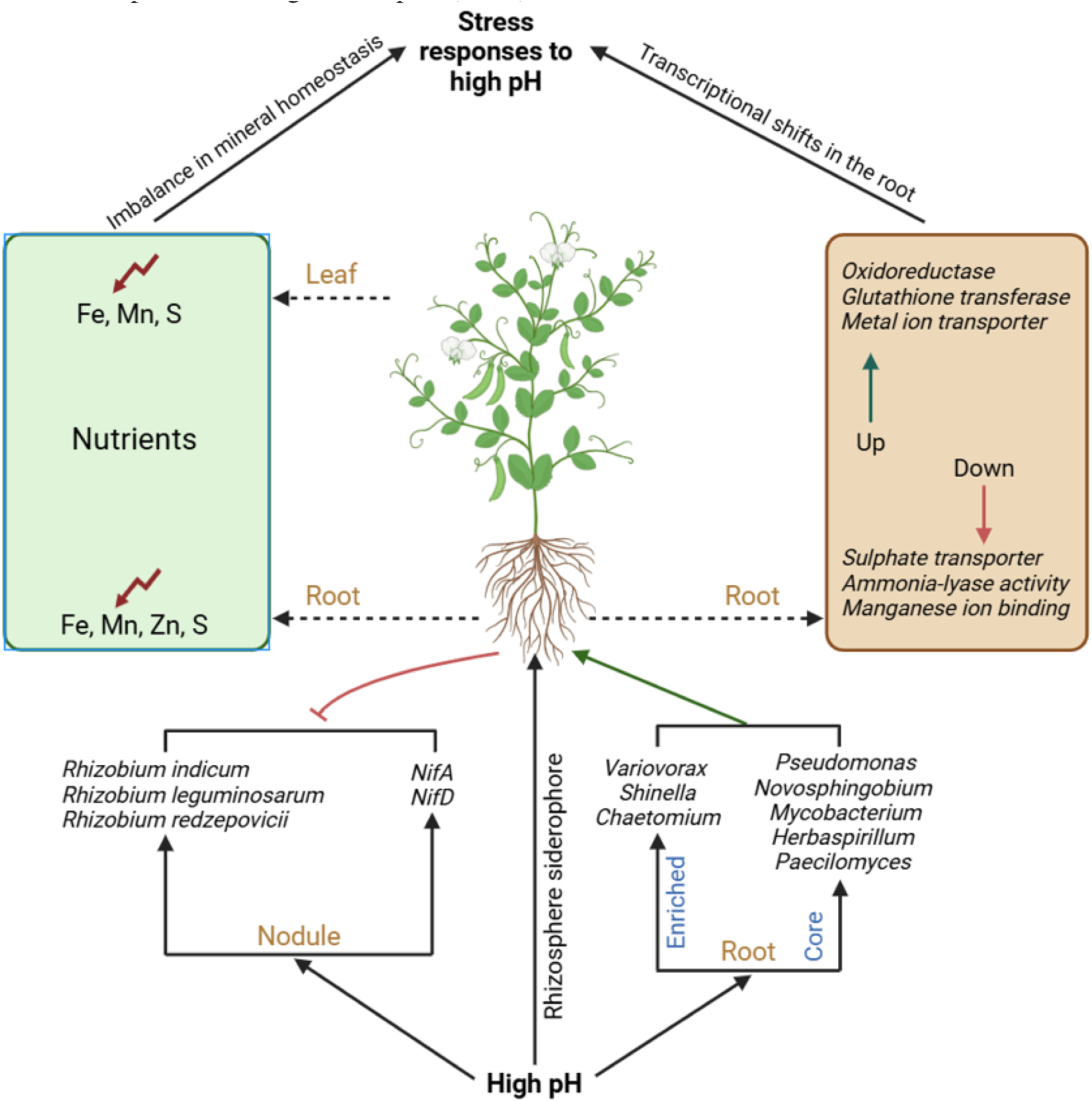
Model of adaptive responses in garden pea under high pH in soil. High pH stress disrupts mineral homeostasis in both roots and leaves, resulting in imbalances in Fe, Mn, Zn, and S levels. Transcriptional shifts in the root include upregulation of oxidative stress response gene (e.g., oxidoreductase, glutathione transferase, and metal ion transporters) and downregulation of genes involved in sulfate transport, ammonia-lyase activity, and manganese ion binding. In nodules, high pH stress alters the abundance of nitrogen-fixing and reduces expression of *NifA* and *NifD* genes, disrupting nitrogen fixation. High pH stress enriches stress-associated microbe such as *Variovorax*, *Shinella*, and *Chaetomium* while maintaining a core microbial community, including genera like *Pseudomonas*, *Novosphingobium*, *Mycobacterium*, *Herbaspirillum*, and *Paecilomyces*.

## Acknowledgments

We express our gratitude to the Genomics Core of Michigan State University and CD Genomics. This research was supported by a startup grant (5SFAES-293007) and Elizabeth and Hayden Cutler’s Endowed Professorship in Biotechnology from the University of Louisiana at Monroe.

## Author contributions

AT cultivated the plants, measured physiological data, extracted RNA, performed 16S amplicon sequencing analysis and prepared the draft manuscript. MRH extracted the DNA and prepared the 16S library for Nanopore sequencing. AHK conceived the idea for the study, interpreted RNA-seq data, supervised the work, and critically revised the manuscript.

## Data availability

Illumina sequencing data was submitted to NCBI under the following BioProject accession numbers: RNA-seq (PRJNA1116995), 16S Amplicon (PRJNA1169338), ITS amplicon (PRJNA1195628) and 16S Nanopore (PRJNA1117024)

## References

1. Acuña, J. J., Rilling, J. I., Inostroza, N. G., Zhang, Q., Wick, L. Y., Sessitsch, A., & Jorquera, M. A. (2024). *Variovorax* sp. strain P1R9 applied individually or as part of bacterial consortia enhances wheat germination under salt stress conditions. Scientific reports, 14(1), 2070.

2. Anders, S., Pyl, P.T., Huber, W., 2015. HTSeq–a Python framework to work with high throughput sequencing data. Bioinformatics 31 (2), 166–169.

3. Assaha, D. V. M., Ueda, A., Saneoka, H., Al-Yahyai, R., & Yaish, M. W. (2017). The Role of Na+ and K^+^ Transporters in Salt Stress Adaptation in Glycophytes. Frontiers in physiology, 8, 509.

4. Aung, T. T., Shi, F., Zhai, Y., Xue, J., Wang, S., Ren, X., & Zhang, X. (2022). Acidic and Alkaline Conditions Affect the Growth of Tree Peony Plants via Altering Photosynthetic Characteristics, Limiting Nutrient Assimilation, and Impairing ROS Balance. International journal of molecular sciences, 23(9), 5094.

5. Bais HP, Weir TL, Perry LG, Gilroy S, Vivanco JM (2006) The role of root exudates in rhizosphere interactions with plants and other organisms. Annu Rev Plant Biol 57(1):233– 266

6. Barrow NJ, Hartemink AE. 2023. The effects of pH on nutrient availability depend on both soils and plants. Plant and Soil 487, 21–37.

7. Bhattacharyya, P. N., & Jha, D. K. (2012). Plant growth-promoting rhizobacteria (PGPR): emergence in agriculture. World journal of microbiology & biotechnology, 28(4), 1327– 1350.

8. Bolger, A.M., Lohse, M., Usadel, B., 2014. Trimmomatic: a flexible trimmer for Illumina Sequence Data. Bioinforma. btu170.

9. Brear, E. M., Day, D. A., & Smith, P. M. (2013). Fe: an essential micronutrient for the legume-rhizobium symbiosis. Frontiers in plant science, 4, 359.

10. Briat, J. F., Dubos, C., & Gaymard, F. (2015). Fe nutrition, biomass production, and plant product quality. Trends in plant science, 20(1), 33–40.

11. Briat, J. F., Dubos, C., & Gaymard, F. (2015). Fe nutrition, biomass production, and plant product quality. Trends in plant science, 20(1), 33–40.

12. Cass, C. L., Peraldi, A., Dowd, P. F., Mottiar, Y., Santoro, N., Karlen, S. D., Bukhman, Y. V., Foster, C. E., Thrower, N., Bruno, L. C., Moskvin, O. V., Johnson, E. T., Willhoit, M. E., Phutane, M., Ralph, J., Mansfield, S. D., Nicholson, P., & Sedbrook, J. C. (2015). Effects of PHENYLALANINE AMMONIA LYASE (PAL) knockdown on cell wall composition, biomass digestibility, and biotic and abiotic stress responses in *Brachypodium*. Journal of Experimental Botany, 66(14), 4317–4335.

13. Chang, J. S., Chou, C. L., Lin, G. H., Sheu, S. Y., & Chen, W. M. (2005). *Pseudoxanthomonas kaohsiungensis*, sp. nov., a novel bacterium isolated from oil-polluted site produces extracellular surface activity. Systematic and applied microbiology, 28(2), 137–144.

14. Chen, W., Koide, R. T., Adams, T. S., DeForest, J. L., Cheng, L., Eissenstat, D. M. (2016). Root morphology and mycorrhizal symbioses together shape nutrient foraging strategies of temperate trees. Proceedings of the National Academy of Sciences of the United States of America, 113(31), 8741–8746.

15. Chieb, M., & Gachomo, E. W. (2023). The role of plant growth promoting rhizobacteria in plant drought stress responses. BMC plant biology, 23(1), 407.

16. Dahl, W. J., Foster, L. M., & Tyler, R. T. (2012). Review of the health benefits of peas (*Pisum sativum* L.). British Journal of Nutrition, 108(S1), S3–S10.

17. Dasila, H., Sah, V. K., Jaggi, V., Kumar, A., Tewari, L., Taj, G., Chaturvedi, S., Perveen, K., Bukhari, N. A., Siang, T. C., & Sahgal, M. (2023). Cold-tolerant phosphate-solubilizing Pseudomonas strains promote wheat growth and yield by improving soil phosphorous (P) nutrition status. Frontiers in microbiology, 14, 1135693.

18. De Vries F. T., Griffiths R. I., Bailey M., Craig H., Girlanda M., Gweon H. S., et al. (2018). Soil bacterial networks are less stable under drought than fungal networks. Nat. Commun. 9:3033.

19. Deng, Y., Kong, W., Zhang, X., Zhu, Y., Xie, T., Chen, M., Zhu, L., Sun, J., Zhang, Z., Chen, C., Zhu, C., Yin, H., Huang, S., & Gu, Y. (2024). Rhizosphere microbial community enrichment processes in healthy and diseased plants: implications of soil properties on biomarkers. Frontiers in microbiology, 15, 1333076.

20. Diby, P.; Park, K.S. Dentification of Volatiles Produced by Cladosporium cladosporioides CL-1, a Fungal Biocontrol Agent That Promotes Plant Growth. Sensors 2013, 13, 13969–13977.

21. Dusha I. (2002). Nitrogen control of bacterial signal production in *Rhizobium meliloti*-alfalfa symbiosis. Indian journal of experimental biology, 40(9), 981–988.

22. Enagbonma, B. J., Fadiji, A. E., Ayangbenro, A. S., & Babalola, O. O. (2023). Communication between Plants and Rhizosphere Microbiome: Exploring the Root Microbiome for Sustainable Agriculture. Microorganisms, 11(8), 2003.

23. Ferrarezi, R. S., Lin, X., Gonzalez Neira, A. C., Tabay Zambon, F., Hu, H., Wang, X., Huang, J. H., & Fan, G. (2022). Substrate pH Influences the Nutrient Absorption and Rhizosphere Microbiome of Huanglongbing-Affected Grapefruit Plants. Frontiers in plant science, 13, 856937.

24. Garnica, M., Bacaicoa, E., Mora, V., San Francisco, S., Baigorri, R., Zamarreño, A. M., & Garcia-Mina, J. M. (2018). Shoot Fe status and auxin are involved in Fe deficiency-induced phytosiderophores release in wheat. BMC Plant Biology, 18(1), 105.

25. Ge, S.X., Jung, D., Yao, R., 2020. ShinyGO: a graphical gene-set enrichment tool for animals and plants. Bioinformatics 36 (8), 2628–2629.

26. Gopalakrishnan, S., Sathya, A., Vijayabharathi, R., Varshney, R. K., Gowda, C. L., & Krishnamurthy, L. (2015). Plant growth promoting rhizobia: challenges and opportunities. 3 Biotech, 5(4), 355–377.

27. Goyal, R. K., Mattoo, A. K., & Schmidt, M. A. (2021). Rhizobial-Host Interactions and Symbiotic Nitrogen Fixation in Legume Crops Toward Agriculture Sustainability. Frontiers in microbiology, 12, 669404.

28. Hamayun, M., Khan, S. A., Khan, A. L., Rehman, G., Kim, Y. H., Iqbal, I., Hussain, J., Sohn, E. Y., & Lee, I. J. (2010). Gibberellin production and plant growth promotion from pure cultures of Cladosporium sp. MH-6 isolated from cucumber (Cucumis sativus L.). Mycologia, 102(5), 989–995.

29. Hasanuzzaman, M., Bhuyan, M. H. M. B., Anee, T. I., Parvin, K., Nahar, K., Mahmud, J. A., & Fujita, M. (2019). Regulation of Ascorbate-Glutathione Pathway in Mitigating Oxidative Damage in Plants under Abiotic Stress. Antioxidants (Basel, Switzerland), 8(9), 384.

30. Hider, R.C., Kong, X., 2010. Chemistry and biology of siderophores. Nat. Prod. Rep. 27 (5), 637–657.

31. Himpsl, S.D., Mobley, H.L.T., 2019. Siderophore Detection Using Chrome Azurol S and cross-feeding assays. Met. Mol. Biol. 2021, 97–108.

32. Hu, K., 2021. Become competent in generating RNA-seq heat maps in one day for novices without prior R experience. Met. Mol. Biol. 2239, 269–303.

33. Ibrahim, S. R. M., Mohamed, S. G. A., Sindi, I. A., & Mohamed, G. A. (2021). Biologically active secondary metabolites and biotechnological applications of species of the family Chaetomiaceae (Sordariales): An updated review from 2016 to 2021. Mycological Progress, 20(595–639).

34. Ibrahim, S. R. M., Mohamed, S. G. A., Sindi, I. A., & Mohamed, G. A. (2021). Biologically active secondary metabolites and biotechnological applications of species of the family Chaetomiaceae (Sordariales): An updated review from 2016 to 2021. Mycological Progress, 20(595–639).

35. Islam S, Akanda AM, Prova A, Islam MdT, Hossain MdM (2015) isolation and identification of plant growth promoting rhizobacteria from cucumber rhizosphere and their effect on plant growth promotion and disease suppression. Front Microbiol 6:1360.

36. Jiang, F., Chen, L., Belimov, A. A., Shaposhnikov, A. I., Gong, F., Meng, X., Hartung, W., Jeschke, D. W., Davies, W. J., & Dodd, I. C. (2012). Multiple impacts of the plant growth-promoting rhizobacterium Variovorax paradoxus 5C-2 on nutrient and ABA relations of Pisum sativum. Journal of experimental botany, 63(18), 6421–6430.

37. Kabir, A. H., & Bennetzen, J. L. (2024). Molecular insights into the mutualism that induces Fe deficiency tolerance in sorghum inoculated with *Trichoderma harzianum*. Microbiological research, 281, 127630.

38. Kabir, A. H., Paltridge, N. G., Able, A. J., Paull, J. G., & Stangoulis, J. C. (2012). Natural variation for Fe-efficiency is associated with upregulation of Strategy I mechanisms and enhanced citrate and ethylene synthesis in Pisum sativum L. Planta, 235(6), 1409–1419.

39. Kabir, A. H., Thapa, A., Hasan, M. R., & Parvej, M. R. (2024). Local signal from *Trichoderma afroharzianum* T22 induces host transcriptome and endophytic microbiome leading to growth promotion in sorghum. Journal of experimental botany, erae340.

40. Kaya, C., Akram, N. A., & Ashraf, M. (2019). Influence of exogenously applied nitric oxide on strawberry (Fragaria × ananassa) plants grown under High pH and/or saline stress. Physiologia Plantarum, 165(2), 247–263.

41. Kim, D. S., & Hwang, B. K. (2014). An important role of the pepper phenylalanine ammonia-lyase gene (PAL1) in salicylic acid-dependent signalling of the defence response to microbial pathogens. Journal of experimental botany, 65(9), 2295–2306.

42. Kim, D., Langmead, B., Salzberg, S.L., 2015. HISAT: a fast spliced aligner with low memory requirements. Nat. Methods 12 (4), 357–360.

43. Kobayashi, T., & Nishizawa, N. K. (2012). Fe uptake, translocation, and regulation in higher plants. Annual review of plant biology, 63, 131–152.

44. Kroh, G. E., & Pilon, M. (2020). Regulation of Fe homeostasis and use in chloroplasts. International journal of molecular sciences, 21(9), 3395.

45. Ku, Y. S., Cheng, S. S., Ng, M. S., Chung, G., & Lam, H. M. (2022). The Tiny Companion Matters: The Important Role of Protons in Active Transports in Plants. International journal of molecular sciences, 23(5), 2824.

46. Küpper, H., Benedikty, Z., Morina, F., Andresen, E., Mishra, A., & Trtílek, M. (2019). Analysis of OJIP chlorophyll fluorescence kinetics and QA reoxidation kinetics by direct fast imaging. Plant Physiology, 179(2), 369–381.

47. Li, N., Liu, H., Sun, J., Zheng, H., Wang, J., Yang, L., Zhao, H., & Zou, D. (2018). Transcriptome analysis of two contrasting rice cultivars during alkaline stress. Scientific reports, 8(1), 9586.

48. Li, P., Teng, C., Zhang, J., Liu, Y., Wu, X., & He, T. (2023). Characterization of drought stress-mitigating Rhizobium from faba bean (*Vicia faba* L.) in the Chinese Qinghai-Tibet Plateau. Frontiers in microbiology, 14, 1212996.

49. Lindström, K., & Mousavi, S. A. (2020). Effectiveness of nitrogen fixation in rhizobia. Microbial biotechnology, 13(5), 1314–1335.

50. Liu, Q., Zhao, X., Liu, Y., Xie, S., Xing, Y., Dao, J., Wei, B., Peng, Y., Duan, W., & Wang, Z. (2021). Response of Sugarcane Rhizosphere Bacterial Community to Drought Stress. Frontiers in microbiology, 12, 716196.

51. Livak, K.J., Schmittgen, T.D., 2001. Analysis of relative gene expression data using realtime quantitative PCR and the 2 ΔΔCT method. Methods 25, 402–408.

52. Luo, J., Liu, T., Diao, F., Hao, B., Zhang, Z., Hou, Y., & Guo, W. (2023). Shift in rhizospheric and endophytic microbial communities of dominant plants around Sunit Alkaline Lake. The Science of the total environment, 867, 161503.

53. Marschner, H., Römheld, V., & Kissel, M. (1986). Different strategies in higher plants in mobilization and uptake of Fe. Journal of Plant Nutrition, 9(3–7), 695–713.

54. Meisrimler, C. N., Wienkoop, S., Lyon, D., Geilfus, C. M., & Lüthje, S. (2016). Long-term High pH: Tracing changes in the proteome of different pea (*Pisum sativum* L.) cultivars. Journal of proteomics, 140, 13–23.

55. Merry, R., Espina, M. J., Lorenz, A. J., & Stupar, R. M. (2022). Development of a controlled-envFement assay to induce Fe deficiency chlorosis in soybean by adjusting calcium carbonates, pH, and nodulation. Plant methods, 18(1), 36.

56. Nakayama, M., Hosoya, K., Tomiyama, D., Tsugukuni, T., Matsuzawa, T., Imanishi, Y., & Yaguchi, T. (2013). Method for rapid detection and identification of *Chaetomium* and evaluation of resistance to peracetic acid. Journal of Food Protection, 76(6), 999–1005.

57. Nouri, M. Z., Moumeni, A., & Komatsu, S. (2015). Abiotic Stresses: Insight into Gene Regulation and Protein Expression in Photosynthetic Pathways of Plants. International journal of molecular sciences, 16(9), 20392–20416.

58. Oláh, B., Kiss, E., Györgypál, Z., Borzi, J., Cinege, G., Csanádi, G., Batut, J., Kondorosi, A., & Dusha, I. (2001). Mutation in the ntrR gene, a member of the vap gene family, increases the symbiotic efficiency of *Sinorhizobium meliloti*. Molecular plant-microbe interactions, 14(7), 887–894.

59. Pinedo, I., Ledger, T., Greve, M., & Poupin, M. J. (2015). *Burkholderia phytofirmans* PsJN induces long-term metabolic and transcriptional changes involved in *Arabidopsis thaliana* salt tolerance. Frontiers in plant science, 6, 466.

60. Qiao, S., Ma, C., Li, H., Zhang, Y., Zhang, M., Zhao, W., & Liu, B. (2024). Responses of growth and photosynthesis to alkaline stress in three willow species. Scientific reports, 14(1), 14672.

61. Răut, I., Călin, M., Capră, L., Gurban, A.-M., Doni, M., Radu, N., & Jecu, L. (2021). Cladosporium sp. isolate as fungal plant growth promoting agent. Agronomy, 11(2), 392.

62. Ravet, K., & Pilon, M. (2013). Copper and Fe homeostasis in plants: the challenges of oxidative stress. Antioxidants & redox signaling, 19(9), 919–932.

63. Ravet, K., & Pilon, M. (2013). Copper and Fe homeostasis in plants: the challenges of oxidative stress. Antioxidants & redox signaling, 19(9), 919–932.

64. Rivas, S., & Thomas, C. M. (2005). Molecular interactions between tomato and the leaf mold pathogen Cladosporium fulvum. Annual review of phytopathology, 43, 395–436.

65. Saeed, Q., Xiukang, W., Haider, F. U., Kučerik, J., Mumtaz, M. Z., Holatko, J., Naseem, M., Kintl, A., Ejaz, M., Naveed, M., Brtnicky, M., & Mustafa, A. (2021). Rhizosphere Bacteria in Plant Growth Promotion, Biocontrol, and Bioremediation of Contaminated Sites: A Comprehensive Review of Effects and Mechanisms. International journal of molecular sciences, 22(19), 10529.

66. Samac, D. A., & Graham, M. A. (2007). Recent advances in legume-microbe interactions: recognition, defense response, and symbiosis from a genomic perspective. Plant Physiology, 144(2), 582–587.

67. Samborska-Skutnik, I. A., Kalaji, H. M., Sieczko, L., & Bąba, W. (2020). Structural and functional response of photosynthetic apparatus of radish plants to High pH. Photosynthetica, 58.

68. Sankari, S., Babu, V. M. P., Bian, K., Alhhazmi, A., Andorfer, M. C., Avalos, D. M., Smith, T. A., Yoon, K., Drennan, C. L., Yaffe, M. B., Lourido, S., & Walker, G. C. (2022). A haem-sequestering plant peptide promotes Fe uptake in symbiotic bacteria. Nature Microbiology, 7(9), 1453–1465.

69. Schwember, A. R., Schulze, J., Del Pozo, A., & Cabeza, R. A. (2019). Regulation of Symbiotic Nitrogen Fixation in Legume Root Nodules. Plants (Basel, Switzerland), 8(9), 333.

70. Shanthiyaa, V., Karthikeyan, G., & Raguchander, T. (2014). Production of extracellular proteins, cellulases, and antifungal metabolites by *Chaetomium globosum* Kunze ex. Fr. Archives of Phytopathology and Plant Protection, 47(5), 517–528.

71. Shee, R., Ghosh, S., Khan, P., Sahid, S., Roy, C., Shee, D., Paul, S., & Datta, R. (2022). Glutathione regulates transcriptional activation of Fe transporters via S-nitrosylation of bHLH factors to modulate subcellular Fe homoeostasis. Plant, cell & Environment, 45(7), 2176–2190.

72. Shumilina, J., Soboleva, A., Abakumov, E., Shtark, O. Y., Zhukov, V. A., & Frolov, A. (2023). Signaling in Legume-Rhizobia Symbiosis. International journal of molecular sciences, 24(24), 17397.

73. Singh, S., Singh, A., Dey, R., Mahatma, M., Reddy, K., Singh, A. K., Gangadhara, K., & Bishi, S. K. (2021). Insights into the physiological and molecular responses of plants to Fe and zinc deficiency. Plant Physiology Reports, 1–10.

74. Sinha, A. K., Jaggi, M., Raghuram, B., & Tuteja, N. (2011). Mitogen-activated protein kinase signaling in plants under abiotic stress. Plant signaling & behavior, 6(2), 196–203.

75. Slatni, T., Vigani, G., Salah, I. Ben, Kouas, S., Dell’Orto, M., Gouia, H., Zocchi, G., & Abdelly, C. (2011). Metabolic changes of Fe uptake in N_2_-fixing common bean nodules during High pH. Plant Science, 181(2), 151–158.

76. Spinelli, V., Brasili, E., Sciubba, F., Ceci, A., Giampaoli, O., Miccheli, A., Pasqua, G., & Persiani, A. M. (2022). Biostimulant Effects of *Chaetomium globosum* and *Minimedusa polyspora* Culture Filtrates on Cichorium intybus Plant: Growth Performance and Metabolomic Traits. Frontiers in plant science, 13, 879076.

77. Sun SL, Yang WL, Fang WW, Zhao YX, Guo L, Dai YJ. The Plant Growth-Promoting *Rhizobacterium Variovorax boronicumulans* CGMCC 4969 Regulates the Level of Indole-3-Acetic Acid Synthesized from Indole-3-Acetonitrile. Appl Environ Microbiol. 2018;84(16):e00298–18.

78. Sun, S. L., Yang, W. L., Fang, W. W., Zhao, Y. X., Guo, L., & Dai, Y. J. (2018). The Plant Growth-Promoting Rhizobacterium *Variovorax boronicumulans* CGMCC 4969 Regulates the Level of Indole-3-Acetic Acid Synthesized from Indole-3-Acetonitrile. Applied and Environmental Microbiology, 84(16), e00298–18.

79. Terry N., Low G. (1982). Leaf chlorophyll content and its relation to the intracellular location of Fe. J. Plant Nutr. 5 301–310.

80. Tewari, R. K., Hadacek, F., Sassmann, S., & Lang, I. (2013). Fe deprivation-induced reactive oxygen species generation leads to non-autolytic PCD in Brassica napus leaves. Environmental and experimental botany, 91(100), 74–83.

81. Thounaojam, N., Sharma, G. D., & Pandey, P. (2018). Composting of rice-residues using lignocellulolytic plant-probiotic *Stenotrophomonas maltophilia*, and its evaluation for growth enhancement of Oryza sativa L. Environmental Sustainability, 1(3), 185–196.

82. Turner AJ, Arzola CI, Nunez GH. (2020). High pH Stress Affects Root Morphology and Nutritional Status of Hydroponically Grown Rhododendron (*Rhododendron* spp.). Plants (Basel). 9(8):1019.

83. Turner, A. J., Arzola, C. I., & Nunez, G. H. (2020). High pH Stress Affects Root Morphology and Nutritional Status of Hydroponically Grown Rhododendron (*Rhododendron* spp.). Plants (Basel, Switzerland), 9(8), 1019.

84. Vigani, G., & Murgia, I. (2018). Fe-Requiring Enzymes in the Spotlight of Oxygen. Trends in plant science, 23(10), 874–882.

85. Wang, Q., Liu, J., & Zhu, H. (2018). Genetic and Molecular Mechanisms Underlying Symbiotic Specificity in Legume-Rhizobium Interactions. Frontiers in plant science, 9, 313.

86. Witte, C. P., & Herde, M. (2020). Nucleotide Metabolism in Plants. Plant Physiology, 182(1), 63–78.

87. Wu, T., Zhang, H. T., Wang, Y., Jia, W. S., Xu, X. F., Zhang, X. Z., & Han, Z. H. (2012). Induction of root Fe(lll) reductase activity and proton extrusion by High pH is mediated by auxin-based systemic signalling in *Malus xiaojinensis*. Journal of Experimental Botany, 63(2), 859–870.

88. Xia, Y., Feng, J., Zhang, H., Xiong, D., Kong, L., Seviour, R., & Kong, Y. (2024). Effects of soil pH on the growth, soil nutrient composition, and rhizosphere microbiome of *Ageratina adenophora*. PeerJ, 12, e17231.

89. Yates, A. D., Allen, J., Amode, R. M., Azov, A. G., Barba, M., Becerra, A., Bhai, J., Campbell, L. I., Carbajo Martinez, M., Chakiachvili, M., Chougule, K., Christensen, M., Contreras-Moreira, B., Cuzick, A., Da Rin Fioretto, L., Davis, P., De Silva, N. H., Diamantakis, S., Dyer, S., Elser, J., … Flicek, P. (2022). Ensembl Genomes 2022: an expanding genome resource for non-vertebrates. Nucleic acids research, 50(D1), D996–D1003.

90. Zakry FAA, Shamsuddin ZH, Rahim KA, Zakaria ZZ, Rahim AA (2012) Inoculation of Bacillus sphaericus UPMB-10 to young oil palm and measurement of its uptake of fixed nitrogen using the 15 N isotope dilution technique. Microbes Environ 27(3):257– 262.

91. Zhang, J. C., Wang, X. N., Sun, W., Wang, X. F., Tong, X. S., Ji, X. L., An, J. P., Zhao, Q., You, C. X., & Hao, Y. J. (2020). Phosphate regulates malate/citrate-mediated Fe uptake and transport in apple. Plant science, 297, 110526.

92. Zhang, W., Chen, Y., Huang, K., Wang, F., & Mei, Z. (2023). Molecular Mechanism and Agricultural Application of the NifA-NifL System for Nitrogen Fixation. International journal of molecular sciences, 24(2), 907.

93. Zhang, X., Zhang, D., Sun, W., & Wang, T. (2019). The Adaptive Mechanism of Plants to Fe Deficiency via Fe Uptake, Transport, and Homeostasis. International journal of molecular sciences, 20(10), 2424.

94. Zhou, Y., Liu, D., Li, F., Dong, Y., Jin, Z., Liao, Y., Li, X., Peng, S., Delgado-Baquerizo, M., & Li, X. (2024). Superiority of native soil core microbiomes in supporting plant growth. Nature communications, 15(1), 6599.

95. Zhu, X. F., Dong, X. Y., Wu, Q., & Shen, R. F. (2019). Ammonium regulates high pH responses by enhancing nitric oxide signaling in *Arabidopsis thaliana*. Planta, 250(4), 1089–1102.

96. Zhuang, L., Li, Y., Wang, Z., Yu, Y., Zhang, N., Yang, C., Zeng, Q., & Wang, Q. (2021). Synthetic community with six Pseudomonas strains screened from garlic rhizosphere microbiome promotes plant growth. Microbial biotechnology, 14(2), 488–502.

