## Supplementary for "Transcriptional reprogramming and microbiome dynamics in garden pea exposed to high pH stress during vegetative stage"

**Supplementary Table S1.** Physico-chemical properties of soil used in the experiments.

| **Texture** | **Organic matter** | **Cation Exchange Capacity** | **Water Holding Capacity** |
|---|---|---|---|
| Clay | 5% | 15.6 meq/100g | 42% |

**Elemental content ppm**

| **Al** aluminum | **As** arsenic | **B** boron | **Ca** calcium | **Cd** cadmium | **Cr** chromium | **Cu** copper | **Fe** iron | **K** potassium | **Mg** magnesium | **Mn** manganese | **Mo** molybdenum | **Na** sodium | **Ni** nickel | **P** phosphorus | **Pb** lead | **S** sulfur | **Zn** zinc |
|---|---|---|---|---|---|---|---|---|---|---|---|---|---|---|---|---|---|
| no limit | <20 | no limit | no limit | <2 | <100 | <100 | no limit | no limit | no limit | <3500 | <440 | no limit | <50 | no limit | <75 | no limit | <100 |
| 7686 | 4.74 | 8.97 | 1415 | 0.24 | 10.2 | 11.0 | 9033 | 1286 | 1831 | 641 | 0.42 | 991 | 9.33 | 626 | 21.0 | 182 | |

**Supplementary Table S2.** Gene-specific primers used in qPCR experiments.

| Gene | Primers |
|---|---|
| *PsGAPDH* | Fw GTGGTCTCCACTGACTTTATTGGT<br>Rv TTCCTGCCTTGGCATCAAA |
| *NifA* | Fw-TGCCGGATGTCTATCCAAAT<br>Rv-AGTCGCAGCGGACAGTAGAT |
| *NifD* | Fw-GGGTCGGATGCATCAAGCAAG<br>Rv-GTTGCTAGGCTCATACGAATATC |
| *NifH* | Fw-CGACCACGTCACAGAACAC<br>Rv-CCTTGTAGCCGACCTTCATG |

**Supplementary Table S3.** The top 10 differentially expressed genes in roots of garden pea exposed to high pH relative to controls. Values marked with an asterisk are significant (> Log2 fold, < P value 0.05).

| **Upregulated genes** | High pH vs Control (fold change) |
|---|---|
| Psat1g015240 (oxidoreductase activity) | 5.1 |
| Psat1g010280 (glutathione transferase activity) | 4.6 |
| Psat1g011640 (metal ion transmembrane transporter) | 3.8 |
| Psat6g220320 (metal ion binding) | 4.4 |
| Psat1g168160 (peroxidase activity) | 4.2 |
| Psat1g002000 (zinc ion binding) | 3.0 |
| Psat5g170360 (Fe ion binding) | 3.2 |
| Psat2g009080 (ABC−type transporter activity) | 2.3 |
| Psat1g096960 (magnesium ion binding) | 2.3 |
| Psat7g119720 (cysteine synthase activity) | 2.9 |
| **Downregulated genes** | |
| Psat1g040440 (sulfate transporter activity) | -2.7 |
| Psat5g270080 (folic acid binding) | -2.1 |
| Psat5g063680 (photosynthesis) | -2.0 |
| Psat5g200040 (chlorophyll binding) | -2.1 |
| Psat1g046920 (ammonia−lyase activity) | -3.0 |
| Psat4g048880 (calmodulin binding) | -2.0 |
| Psat5g094200 (manganese ion binding) | -2.0 |

| | |
|---|---|
| Psat6g173720 (heme binding) | -2.9 |
| Psat1g185960 (glutamate−ammonia ligase activity) | -2.1 |
| Psat2g053680 (monooxygenase activity) | -2.0 |
